# Revisiting the role of microtubules in cytotoxic T cell function: efficient search and killing without microtubule network

**DOI:** 10.1101/2025.02.24.639827

**Authors:** Galia Montalvo, Reza Shaebani, Lukas Schuster, Shweta Nandakumar, Natasha Cowley, Sijia Zhang, Renping Zhao, Anna Burgstaller, Shulagna Sharma, Oskar Staufer, Rhoda Hawkins, Markus Hoth, Marcel A. Lauterbach, Laura Schaedel, Bin Qu, Franziska Lautenschläger

**Affiliations:** Department of Experimental Physics, Saarland University, Saarbrücken, Germany; Biophysics, Center for Integrative Physiology and Molecular Medicine (CIPMM), School of Medicine, Saarland University, Homburg, Germany; Department of Theoretical Physics, Saarland University, Saarbrücken, Germany; Center for Biophysics, Saarland University, Saarbrücken, Germany; Department of Physics and Astronomy, University of Sheffield, Sheffield, United Kingdom; Department of Mathematics, University of Manchester, Manchester, United Kingdom; Department of Biomedical Sciences, Institute for Health Research and Education (IGB), Osnabrück University, Osnabrück, Germany; INM - Leibniz Institute for New Materials, Saarbrücken, Germany; Max Planck School, Matter to Life, Heidelberg, Germany; Molecular Imaging, Center for Integrative Physiology and Molecular Medicine (CIPMM), School of Medicine, Saarland University, Homburg, Germany

**Keywords:** CTL, pretubulysin, Search, Microtubules, Migration

## Abstract

To clean tissue from tumorigenic and infected cells, cytotoxic T lymphocytes (CTLs) must navigate confined environments in vivo, locate the infected cells and eliminate them. Impaired CTL migration towards the tumor can limit the efficacy of immunotherapy. Microtubules (MTs) have emerged as promising targets, because destabilizing MTs enhances T-cell migration and subsequent killing, yet the underlying mechanisms are poorly understood. Here, we use pretubulysin, a potent MT depolymerizer, to uncover how MT dynamics regulate CTL motility. Complete MT disassembly markedly increased CTL infiltration and migration in 3D matrices. To investigate how altered migration affects target elimination, we employed a persistent random-walk model parameterized solely with experimental motility data. The model shows that the increase in speed and persistence induced by pretubulysin explains enhanced search efficiency increasing the encounter rate of CTLs with target cells. The simulations also predict how these gains in search efficiency scale with tissue thickness and CTL density. Mechanistically, MT depolymerization in activated CTLs triggers localized actomyosin accumulation at the uropod. This enhances rear contraction forces and promotes faster, more persistent migration and efficient search. Our findings clarify how MT dynamics influence CTL ability to eliminate targets in 3D environments and highlights the potential of MT-targeting agents such as pretubulysin to optimize T cell-based immunotherapies.

**Significance Statement:** Cytotoxic T lymphocytes (CTLs) must efficiently navigate dense tissues to locate and eliminate target cells, yet their movement is strongly limited by physical constraints. We demonstrate that disrupting the microtubule network with pretubulysin enhances CTL motility by increasing migration speed and persistence, enabling faster infiltration into 3D matrices. Integrating experimental measurements with parameter-free persistent-random-walk simulations reveals that these motility changes alone explain the improved search efficiency. Mechanistically, microtubule disassembly induces a marked reorganization of actomyosin toward the uropod, strengthening rearward contractility and promoting persistent migration. Our findings reveal an important mechanism by which cytoskeletal dynamics influence CTL function. Furthermore, they highlight the potential of MT-disassembling agents to strengthen T-cell-mediated cytotoxicity in future immunotherapeutic strategies.

## Introduction

Cell migration is a key feature of immune function, and immune cells are particularly specialized to migrate in nearly all tissues within the human body (1). Motility and the capability to infiltrate into tissues are essential for cytotoxic T lymphocytes (CTLs) to efficiently locate tumorigenic or pathogen-infected cells and to eliminate them (2). This remarkable capacity arises from optimized migration patterns and search strategies that enhance target detection. Cells can control their search efficiency by tuning migratory properties such as speed and persistence (3–6). We previously described a universal coupling between speed and persistence, in which faster cells move more persistently (7), optimizing the time required to find a target (4, 5). The coupling between speed and persistence depends on both actin polymerization and cell polarization, which coexist inherently in actomyosin-driven motion (7). T cells use amoeboid locomotion, in which cell shape changes dynamically. Cellular polarization, characterized by the rapid generation of protrusions at the leading edge and a uropod at the posterior region, along with high actomyosin contractility, serves as major force-generating machinery (8).

Two major components of the cellular cytoskeleton, actin and microtubules (MT) are dynamically coupled and regulated to coordinate migration (9). During T cell migration, the formation of protrusions at the leading edge is driven by polymerization of branched filamentous actin (F-actin), which pushes the plasma membrane providing the primary driving force for forward movement. Meanwhile, the microtubule-organizing center (MTOC) is located at the uropod, behind the nucleus (10). In contrast, for mesenchymal motility, the MTOC is positioned between the nucleus and the leading edge (11). MTOC positioning plays a decisive role in defining T cell polarity to govern migration direction (10). For migrating T cells, actomyosin contractions at the uropod allow detachment from the substrate for further migration and provide rearward squeezing forces to facilitate movement of the nucleus through confined spaces (8). The seemingly distinctively located networks, actin and MT, can interact and influence each other’s dynamics, including at the interface between T cells and target cells where actomyosin dynamics influence microtubule disassembly (12).

Polarization of MTOC has long been considered a key mechanism for directing lytic granules to the immunological synapse formed between CTLs and their target cells (13). In the prevailing model, lytic granules are transported along microtubules toward the polarized MTOC and are subsequently delivered and secreted at the synaptic interface). However, evidence challenging this paradigm has emerged. For instance, lytic granule release can occur before MTOC polarization in some CTL-target cell conjugates (14) and partial disruption of the MT network with nocodazole does not impair lytic granule secretion or target cell killing (15). Despite these observations, whether an intact MT network is absolutely required for lytic granule delivery and CTL-mediated cytotoxicity has remained unresolved.

*In vivo*, CTLs must navigate through confined 3D environments to locate target cells. The extracellular matrix (ECM), composed of fibrous proteins such as collagen, plays a crucial role in maintaining tissue architecture. During CTL migration in a 3D context, CTLs preferably enter pre-existing confined tunnels in collagen matrices (16). In the context of solid tumors, the ECM often becomes condensed, creating a physical barrier that hinders CTLs infiltration resulting in evasion of immune surveillance (17). This limitation contributes significantly to the low efficacy of adoptive immunotherapy against solid tumors, as confined spaces within the dense ECM impair CTL migration and reduce killing efficiency (15). Recent research has highlighted the potential of targeting MTs to enhance CTL function, particularly in condensed 3D matrices. Disrupting the MT network using agents like nocodazole, or the chemotherapeutic drug Vinblastine significantly improves CTL migration and killing efficiency in 3D environments, especially within dense collagen matrices (15, 18). In T cells, MT network disruption enhances surface tension (18) and activates Rho A (19), a key regulator of cell contractility. Additionally, at the interface between T cells and target cells, MT dynamics play an essential role in regulating cell contractility (20). Similarly, in migrating dendritic cells, MT dynamics influence the protrusion retraction and overall migration, which also involves Rho A (21). Despite these insights, the precise mechanisms by which MT disruption improves T cell motility in 3D environments is not yet fully understood.

Here, using a novel MT-disrupting agent pretubulysin, which efficiently disassembles MT filaments, we demonstrate that with a loss of MTs, CTLs maintain their killing capacity and migrate much faster and consistent in 3D. MT disassembly induces actomyosin redistribution from the leading edge to the uropod, accelerating T cell movement and improving killing efficiency in 3D environments.

## Results

### Integrity of microtubule network negatively regulates search and cytotoxic efficiency of CTLs in dense 3D environments

To manipulate MT dynamics, we used pretubulysin (abbreviated Ptub in all figures) and nocodazole, a well-known and widely used microtubule-disrupting drug (15, 18). Pretubulysin is an innovative drug with potent microtubule depolymerization capacity that is already used to treat cancer cells (22–25). We observed that pretubulysin treatment completely disassembles MT network in CTLs (SI Appendix, Fig. S1). Notably, pretubulysin-induced MT-disassembly could persist for at least 12 hours after washout, consistent with previous reports (26) (SI Appendix, Fig. S1). Together, pretubulysin and nocodazole provide a good model for integrity of MT at different levels.

To examine possible effects of MT integrity on CTL functionality, first we focused on CTL infiltration into 3D matrices. Building on the 3D infiltration assay previously applied to NK cells (27), CTLs were fluorescently labeled with carboxyfluorescein succinimidyl ester (CFSE) and placed on top of collagen matrices, allowing them to infiltrate into a 3D environment (Figure 1A, left). Collagen concentrations of 2 mg/ml and 4 mg/ml were used to mimic healthy tissue and tumor tissue, respectively (28). The focal plane was positioned at the bottom of the matrix, allowing visualization of infiltrated CTLs as they approached the bottom (Movie S1). We found that increasing concentrations of pretubulysin resulted in a greater number of CTLs appearing at the focal plane for both concentrations, especially prominent for dense collagen (4 mg/ml) at 12 hours (Figure 1 B, Movie S1, SI Appendix, Fig. S2). The total number of cells reaching the focal plane after 24 hours remained unaltered between pretubulysin-treated and control CTLs, indicating that the influence of MTs is most pronounced during the early stages of infiltration (Figure 1 B, Movie S1 & SI Appendix, Fig. S2). These findings suggest that integrity of MT negatively regulates CTL infiltration capability.

**Figure 1.**
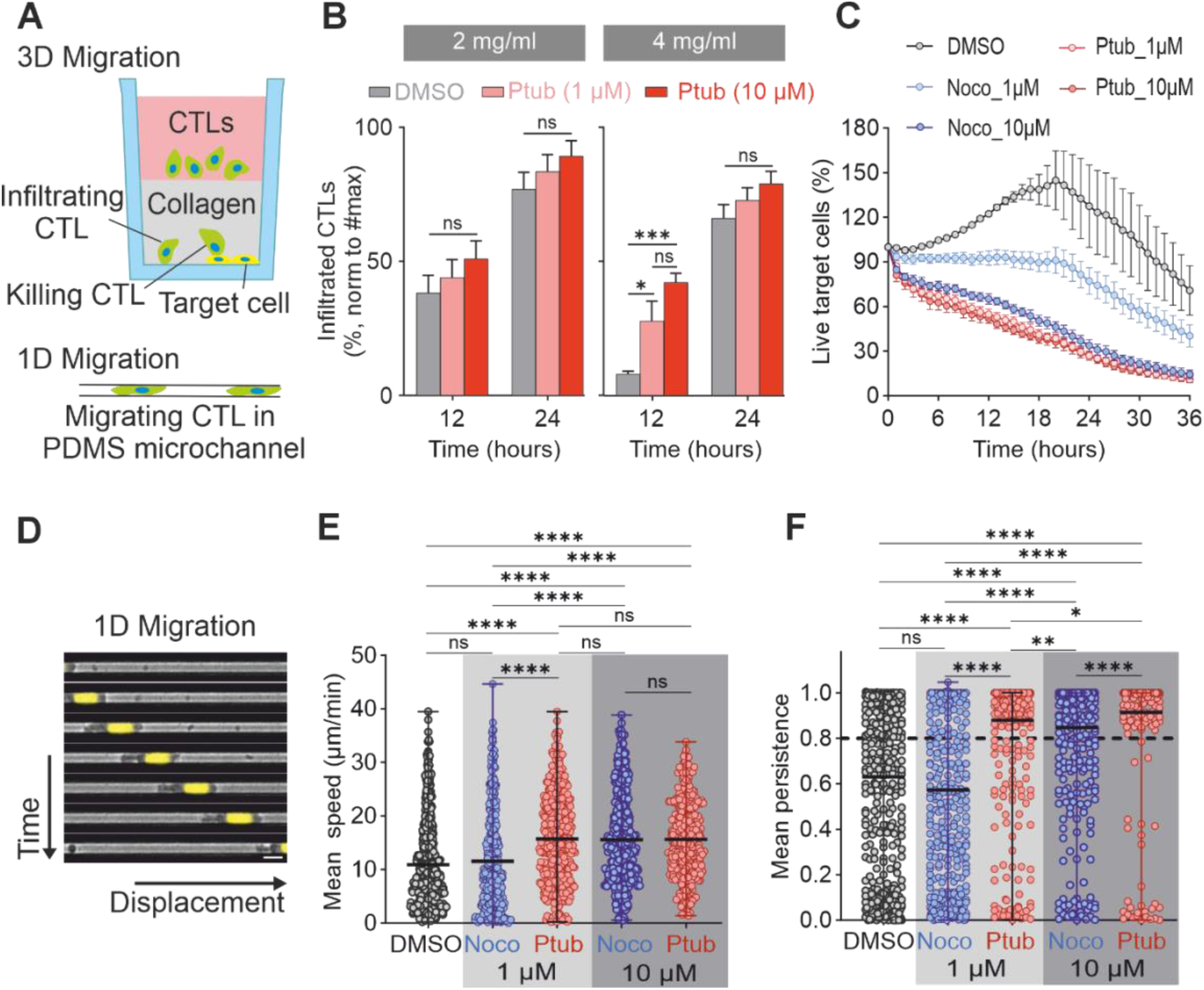
Pretubulysin enhances CTL function in 3D by improving migration parameters. (A) Scheme showing the experimental set up to test CTLs infiltration, killing, and 1D migration. For the infiltration assay CTLs were stained with carboxyfluorescein succinimidyl ester (CFSE), treated with DMSO or pretubulysin (1µM and 10 μM) for 30 min and added on top of solidified collagen matrix (collagen represented in grey, CTLs culture medium without any drug is represented in pink). Cells were visualized as they reached the bottom of the well. In the killing assay tumor cells, expressing pCasper were used as target cells on the bottom of the collagen gel. Live target cells are orange–yellow, apoptotic target cells are green, and dead cells lose fluorescence. Treated CTLs (DMSO, nocodazole or pretubulysin) were placed on top of the collagen gel. 1D migration was performed using PEG-coated PDMS microchannels (length: 400 μm; width: 5 μm; height: 5 μm). (B) Quantification of CTL infiltration at 12 and 24 h in 2 and 4 mg/mL collagen matrices. Bars represent the mean ± SEM of three independent donors (n = 3). For each collagen density, data were analysed using ordinary two-way ANOVA, with treatment and time as factors, followed by Tukey’s multiple-comparisons test to compare DMSO, 1 µM Ptub, and 10 µM Ptub within each time point. Statistical significance is indicated as *p < 0.05, **p < 0.01, and ***p < 0.001. (C) Quantification of target cell death during the 3D killing assay. Dots represent the mean value of four donors, error bars represent the standard deviation (SD). (D) Images taken at different timepoints during the 1D migration assay from one representative donor. Hoechst-stained CTLs (yellow nuclei) were treated with DMSO or drugs and loaded on channels. Cells migrated spontaneously for 15 hours and images were acquired every 2 minutes. Scale bar is 10 μm. (E-F) CTLs, obtained from six donors for DMSO and 1µM treatments and three donors for the 10 µM treatments, were tracked from time lapse videos. In the graphs, each dot represents the mean speed (E) or mean persistence (F) of the track. For statistical analysis, Kruskal-Wallis test and the Dunn’s multiple comparison test were used (* p < 0.05, ** p < 0.01, **** p < 0.0001).

As MTs are believed to play an essential role in CTL killing function, we postulated that a complete MT-disassembly would inhibit CTL-mediated cytotoxicity despite of enhanced infiltration. To test this, we used a 3D killing assay (28) in which tumor cells stably expressing pCasper, an apoptosis reporter, were embedded in dense collagen matrices, and after solidification, CTLs were added from above (Figure 1A, right). We used a ratio of CTLs to tumor cells of 5:1. Time lapse (Movie S2) and subsequent quantifications of the 3D killing assay (Figure 1C & SI Appendix, Fig. S3) show that nocodazole enhanced CTLs killing efficiency in 3D matrices, consistent with previous reports (15, 18). Unexpectedly, pretubulysin-treated CTLs also exhibited a substantially enhanced killing capacity (Figure 1C & SI Appendix, Fig. S3). Remarkably, at the low concentration (1 µM) pretubulysin much more potently enhanced CTL killing efficiency in the 3D matrix relative to nocodazole (Figure 1C & SI Appendix, Fig. S3). Importantly, the effect of pretubulysin at 1 µM is comparable to that of nocodazole at 10 µM (Figure 1C). An increase in the number of target cells in the control treatment was also observed (Figure 1C & SI Appendix, Fig. S3, DMSO), as previously reported (28), indicating that 3D conditions supported target cell proliferation. These data indicate that even in the absence of an MT network, CTLs can still efficiently eliminate target cells.

To understand the underlying mechanism, we examined whether pretubulysin treatment potentiates release of lytic granules using a CD107a degranulation assay. CD107a is a lysosome-associated membrane protein, which becomes incorporated into the plasma membrane upon lytic granule fusion. Our results showed that lytic granule release was not affected in CTLs treated with pretubulysin or nocodazole (SI Appendix, Fig. S4), suggesting that cytotoxic activity is still intact in treated CTLs. The release of lytic granules was not altered and could therefore not explain the increase in killing which we had observed. We hypothesized that cells without an intact MT network might be more efficient in searching and finding the target cells, which would automatically result in higher killing rates. We have shown before that either the speed of migration, the persistence of single trajectories or the coupling between those two parameters decreases the first passage time (4), resulting in more efficient search and would therefore be able to explain the increased killing. We therefore next examined CTL migration using microfabricated channels to mimic the tunnels found in collagen (Figure 1A, D). We observed that disruption of MT network substantially enhanced CTL migration, both in speed and persistence under this confined condition (Figure 1E-F & SI Appendix, Fig. S5). In particular, CTLs with MT disassembly (pretubulysin-treated) were substantially faster (Figure 1E & SI Appendix, Fig. S5A) and more persistent (Figure 1F & SI Appendix, Fig. S5B) compared to the nocodazole-treated groups with partial MT disassembly. Concerning the fraction of highly persistent CTLs (persistence > 0.8), at the low concentration this fraction was doubled for the pretubulysin-treated group (83%) relative to the nocodazole-treated counterparts (40%). At the high concentration, the difference between pretubulysin- and nocodazole-treated CTLs was reduced, but still present (91% vs 74%).

Collectively, these findings demonstrate that the integrity of MT network negatively regulates CTL infiltration by enhancing migration and the consequent killing efficiency in confined environments. Thus, this approach offers a reliable and powerful way to enhance CTLs killing efficiency in a matrix.

### MT disassembly enhances search efficiency under confined conditions, as demonstrated by predictive simulations

To determine whether the experimentally observed increase in target elimination by pretubulysin-treated CTLs could be explained solely by changes in migration behavior, we performed persistent random walk search simulations using only experimentally measured parameters (i.e. speed, persistence, and system geometry); no quantities were adjusted to fit the killing data. CTLs were modeled as persistent random walkers (PRWs) entering the 3D space from the top surface (Figure 2A inset) and migrating until reaching and eliminating target cells at the bottom plane. This framework has previously been shown to accurately reproduce CTL motility in 3D collagen matrices (16). Experimental speed and persistence values for control CTLs served as baseline parameters. For pretubulysin-treated CTLs, we applied the fold-change in speed and persistence observed in microchannel experiments, assuming that migration through collagen channels resembles motion in narrow conduits. This assumption is validated by the close, agreement between parameter-free simulations and experimental killing kinetics (Figure 2A), demonstrating that the enhanced elimination of target cells in pretubulysin-treated samples can be quantitatively accounted for by altered migration dynamics alone. Figure 2A compares the covered area by target cells over time for control and treated cells in experiments and simulations. In the experiments, the covered area is defined as the fraction of the target plane in which fluorescence remains. The bottom plate is initially coated with fluorescent target cells; as CTLs arrive and eliminate these targets, the corresponding regions lose fluorescence. These darkened areas are segmented and quantified, and their cumulative size is subtracted from the total initial fluorescent area. The remaining fluorescent area is then expressed as a percentage. In the simulations, we mimic this process by representing the target layer as a discrete 2D grid that is initially fully occupied (see section Persistent random search simulations in SI Appendix, Materials and Methods for details). When a simulated CTL reaches the bottom plane, it clears a region around its point of impact corresponding to the approximate size of a target cell. The fraction of remaining occupied grid sites over time yields the simulated covered area, directly analogous to the experimentally measured remaining fluorescent area. The quantitative agreement between simulated PRW and experimental data is remarkable since, unlike the role of speed, the influence of persistence on the search efficiency is nontrivial: The mean first-passage time of a persistent random walker in confined geometries is a nonlinear function of the persistence. It even exhibits a nonmonotonic behavior and a confinement-size-dependent optimization upon varying the persistence (4, 29). Thus, an increased persistence of CTLs does not necessarily lead to an improved search efficiency. This means that numerical evaluation is necessary to predict the search time under observed experimental conditions.

**Figure 2.**
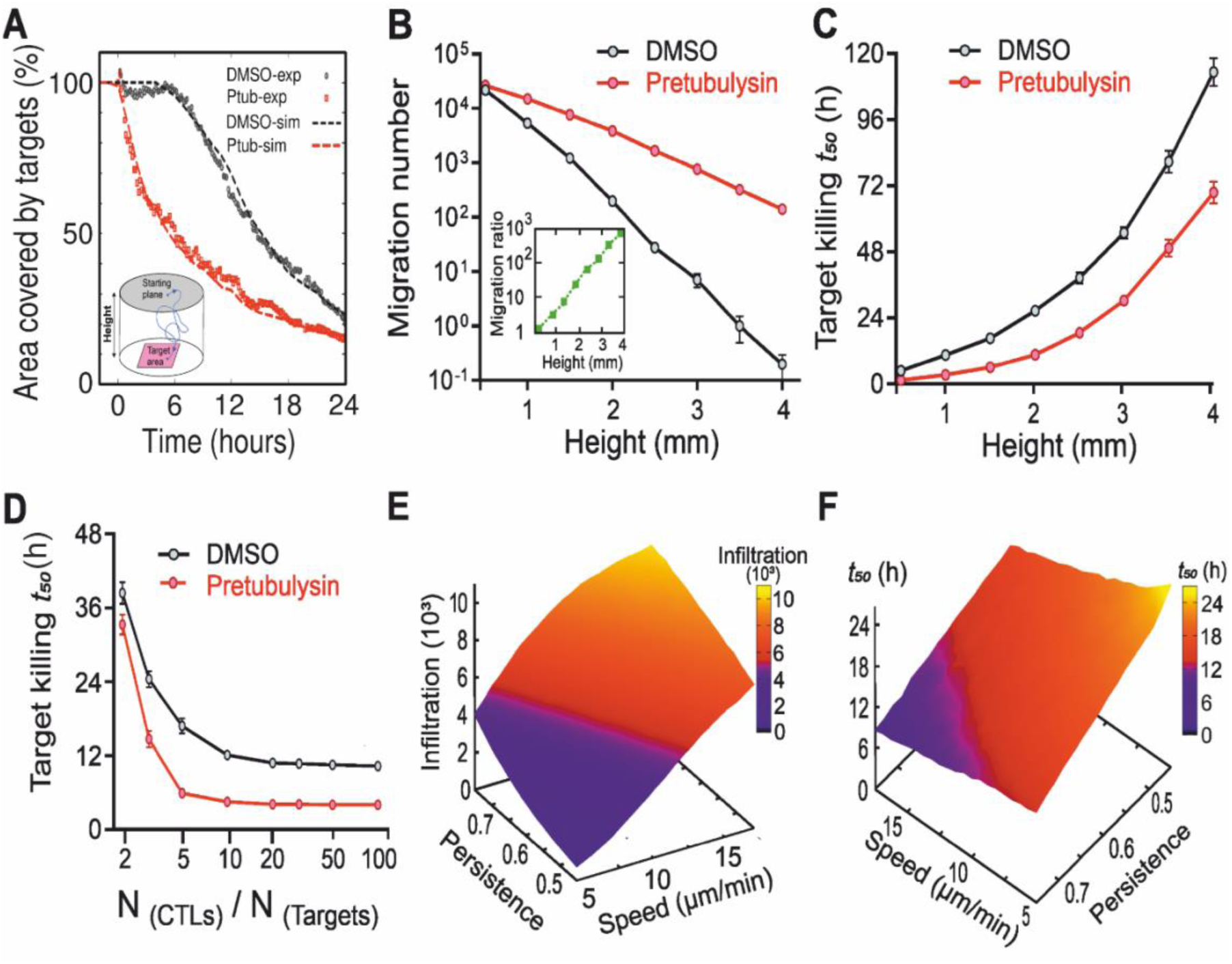
Quantification of migration and killing efficiency of control and pretubulysin-treated CTLs in 3D obtained from persistent random search simulations. (A) Comparison of killing kinetics between simulations and experimental data. Time evolution of the covered area by target cells is shown for experiments and corresponding simulations using the experimental cell motility data. Inset: Schematic of the simulation setup. CTLs were modeled as persistent random walkers entering the medium from the top surface, indicated as “starting plane”, and migrating until they reach the target cells at the bottom plate, indicated as “target area”. (B) Dependence of migration efficiency on the thickness of collagen matrix (height) for control and pretubulysin-treated cells. The inset indicates that the ratio between the number of treated and control cells which reached the target area increases by several orders of magnitude upon a few-fold increase of the collagen matrix height. (C) Killing capacity versus collagen matrix height for treated and control CTLs. The difference between the killing halftime t_50_ (i.e. the time spent to eliminate 50% of the targets) of control and treated cells grows with increasing collagen thickness. (D) Killing capacity in terms of the ratio between the initial number of CTLs and target cells (N_CTLs_/N_targets_) for treated and control CTLs. (E) Number of migrating cells reaching the target area in terms of mean speed and persistence. (F) Killing halftime (t_50_) versus mean speed and persistence.

#### Dependence on matrix thickness

Because CTLs must patrol tissues of varying thickness, we next investigated how system height h affects infiltration. Increasing h reduced the number of CTLs reaching the bottom plane for both groups, as expected for diffusive and persistent search processes. Interestingly, the rate of decrease differed between control and pretubulysin-treated CTLs, resulting in an advantage for the treated group at larger h (Figure 2B). For example, doubling the height from 1.5 to 3.0 mm increased the ratio of arriving treated to control cells from roughly 6 to 108, revealing a highly non-linear amplification of the migration advantage in thicker environments. This trend is also reflected in the killing halftime t_50_, defined as the time required to eliminate 50% of targets. The difference in t_50_ between treated and control CTLs increases with h (Figure 2C), indicating that the functional effect of enhanced migration becomes more pronounced as the search space expands.

#### Dependence on CTL-target ratio

We next varied the initial CTL/target number ratio (N_CTL / N_target) to examine how collective search dynamics affect killing efficiency. For both groups, increasing the number of CTLs accelerated killing and reduced t_50_, but in a nonlinear fashion: the benefit saturated at ratios between 10 and 20 (Figure 2D). This saturation arises from geometric constraints and redundancy in search trajectories and therefore does not follow directly from the immediate-killing assumption. Importantly, across all ratios, pretubulysin-treated CTLs consistently achieved lower t_50_ values than control cells. This effect is even more pronounced at larger ratios but eventually reaches a ratio-independent regime. These results allow two model-based predictions: (i) killing efficiency in a finite 3D domain has an upper limit determined by motility and geometry, and (ii) pretubulysin shifts both the saturation level and the approach to it, but the relative advantage plateaus when CTL density becomes sufficiently large.

#### Combined influence of speed and persistence

Finally, we systematically varied migration speed and persistence within experimentally relevant ranges to assess their combined influence on infiltration and killing. The number of CTLs reaching the target area decreased sharply with reduced speed or persistence (Figure 2E), and the killing halftime increased correspondingly (Figure 2F). We note that the numerical results indicate nonlinear dependencies on the set of motility parameters. While the influence of speed on search performance is intuitive, the dependence on persistence is less so, as we explained above.

### Pretubulysin induces instantaneous MT depolymerization *in vitro* and increases cell stiffness on CTLs

We have confirmed that MT destabilization improves CTLs infiltration, yet we found differences in the effect of the different MT depolymerizing agents pretubulysin and nocodazole. To characterize and compare the MT depolymerizing potential of both drugs at the single, dynamic MT level in a controlled environment, we performed microfluidics-based *in vitro* reconstitution assays using purified tubulin. Briefly, short, biotinylated MT fragments (‘seeds’ in red, Figure 3A, Step I) were immobilized on passivated coverslips via neutravidin. Dynamic MTs (in green, Figure 3A, Step I) were elongated from these seeds using 10 µM purified tubulin (20% ATTO-488-labeled). Subsequently, a drug mix containing pretubulysin/nocodazole (0.1, 1 and 10 µM), 1 mM GTP and 10 µM ATTO-488-labeled tubulin (included to prevent MT disassembly from dilution) was introduced through the microfluidic setup, allowing for live imaging of microtubules during drug treatment. Pretubulysin induced rapid MT depolymerization and resulted in a 60-88% (Figure 3 B-D) reduction in microtubule mass when compared to the control (DMSO). In contrast, nocodazole treatment caused most MTs to enter a paused state characterized by suppressed MT dynamics with no significant shrinkage (SI Appendix, Fig. S6A-E). This behavior is consistent with the observations reported by Vasquez *et al*., 1997 (30). We observed this effect at nocodazole concentrations up to 50 µM. At higher concentrations, MT mass was reduced by only 10.2% (SI Appendix, Fig. S6F). Collectively, these *in vitro* experiments demonstrate that Pretubulysin is a potent MT-depolymerizing agent that induces rapid MT disassembly. In contrast, nocodazole primarily suppresses MT dynamicity whilst causing relatively minimal loss of MT mass.

**Figure 3.**
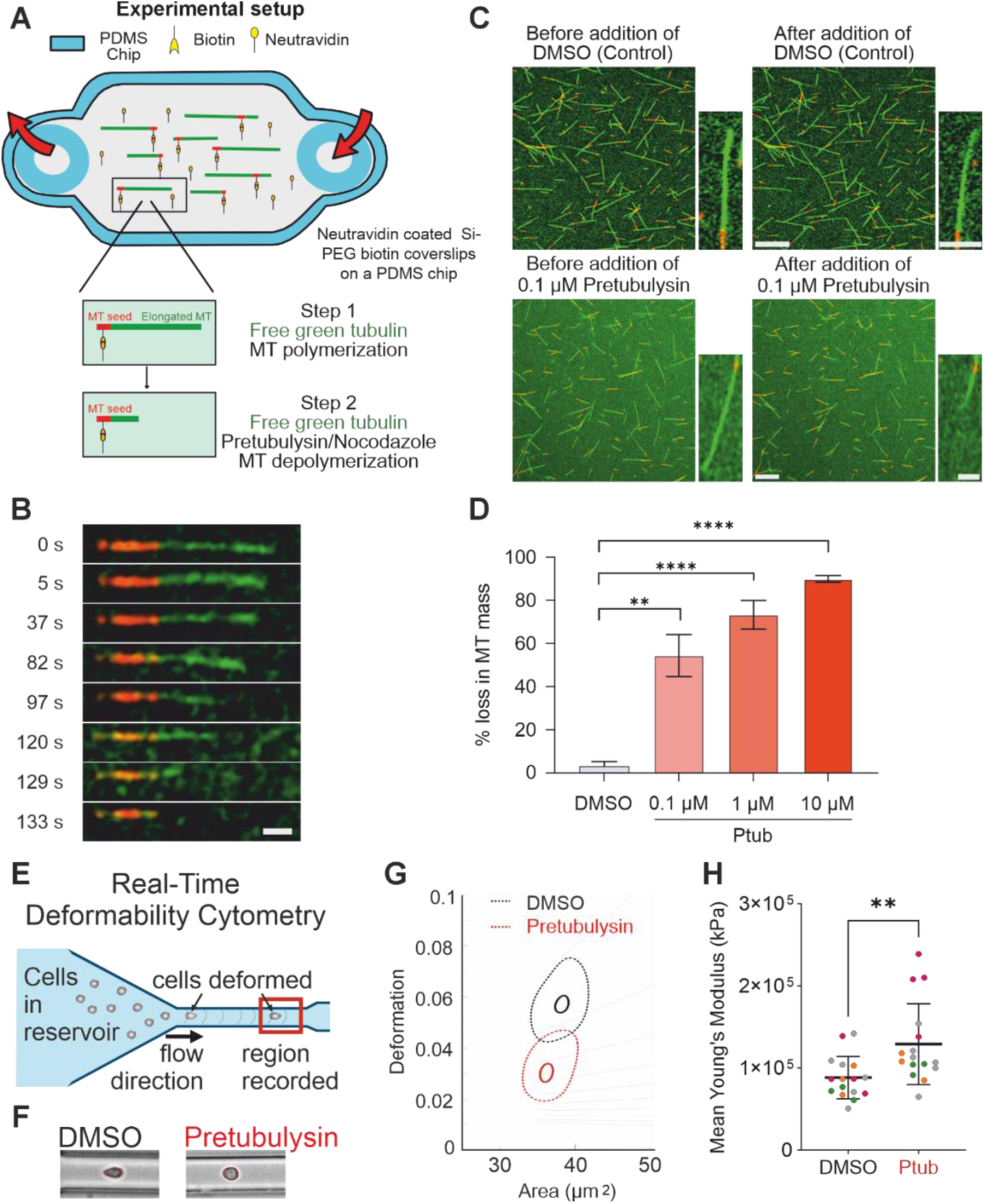
Pretubulysin is a potent microtubule depolymerization agent and stiffens CTLs. (A) Schematic representation of the experimental set-up used to perform microfluidics-based *in vitro* reconstitution assays using purified tubulin: short biotin-containing microtubule seeds (in red) were attached to a neutravidin-coated Si-PEG Biotin passivated coverslip on top of the PDMS chip in a microfluidic circuit. Microtubules (in green) were elongated from these seeds by addition of 10 μM labeled tubulin (Step I). Subsequently the drug mix containing various concentrations of the drug was flushed in (Step II). Images were acquired every 1 s following addition of the drug. (B) Timelapse sequence showing the shrinkage in microtubule length following addition of 10 µM pretubulysin (Scale bar: 2 μm). Images are representative of three independent experiments. (C) Representative image of a field-of-view before and after addition of DMSO (Control, top) and 0.1 µM pretubulysin (bottom) (Scale bar: 5 µm). Inset images show example of a single MT before and after addition (Scale bar: 2 µm). Images are representative of three independent experiments. (D) Comparison of % loss in MT mass following treatment with 0.1, 1 and 10 µM pretubulysin with Control (DMSO). Data represent Mean ± SD from three independent experiments. Unpaired t-test was used for statistical significance (** p < 0.01, **** p<0.0001). (E) Scheme of real-time deformability cytometer setup. A 20 µm microfluidic PDMS chip was assembled on the stage of an inverted microscope. The cell suspension was loaded on the reservoir and deformed by shear stresses and pressure gradient caused by the flow profile. (F) Representative images of cells acquired during real-time deformability cytometry and showing the typical bullet shape for control cells (DMSO) versus round shape observed for pretubulysin-treated CTLs. (G) Representative kernel density estimate plot depicting cell area versus deformation showing pretubulysin treatment makes CTLs less deformable than DMSO-treated control cells. (H) Apparent Younǵs modulus was calculated and analyzed using linear mixed models available with the ShapeOut2.0 software. The results obtained from four different donors are represented on the graph with a color code for each individual donor, where one dot represents one experiment and error bars represent standard deviation of the mean. At least 3,000 events were acquired for each condition in every experiment. For statistical analysis, a paired t-test was used (** p < 0.01).

After having confirmed that pretubulysin efficiently destabilizes MTs and enhances CTLs navigation in confined 3D environments, we sought to understand the underlying mechanism. As MTs are the most rigid cytoskeletal filaments (31), we postulate that MT disassembly induced by pretubulysin softens CTLs, facilitating their infiltration and migration. To test this, we determined the stiffness of CTLs with Real-Time Deformability-Cytometry (RT-DC, Figure 3E). For this, cells were passed through a microfluidic channel (20 µm), and the shear stress-induced cell deformation was used to calculate the apparent Younǵs modulus (32). Pretubulysin treatment reduced deformation without changing cell size (Figure 3F, G). Concomitantly, the apparent Young’s moduli of CTLs were enhanced after pretubulysin treatment compared to that of DMSO-treated counterparts (Figure 3H). These results indicate that MT disassembly not only enhances local surface tension, as previously reported (18), but also leads to overall cell stiffening, reflecting the combined mechanical contributions of the actin cortex and deeper cytoskeletal structures.

### Microtubule disassembly-induced actomyosin enrichment in the uropod increases migration speed

To understand the underlying mechanism, we investigated the role of actin filaments and myosin motors in CTLs with depolymerized MTs. The MT network and dynamics play a critical role in actin cytoskeletal dynamics as well as actomyosin contractility, which are essential for cell motility (33). To further understand how MT disassembly enhances CTLs infiltration and migration in 3D, we examined the distribution of F-actin and phosphorylated myosin (pMyosin) using immunostaining. The actin-associated motor protein myosin II is activated through phosphorylation of its regulatory light chain by upstream signaling pathways involving RhoA, Rho-associated protein kinase (ROCK) and myosin light chain kinase (34, 35). Phosphorylation of myosin II enhances actomyosin contractility, leading to contraction of the cell body and changes in mechanical properties (35–37). Thus, the level and spatial distribution of pMyosin serves as an indicator for the contractile activity of the actomyosin machinery. In the DMSO-treated control CTLs, F-actin was primarily located at the leading edge and around the nucleus, with pMyosin surrounding the nucleus and present in both the protrusions and the uropod (Figure 4A, DMSO). Colocalization of F-actin and pMyosin was observed mainly in the perinuclear region as well as along the contour of the leading edge and the uropod (Figure 4A, DMSO). In comparison, in pretubulysin-treated CTLs, F-actin was primarily located in the uropod and around the nucleus, with pMyosin also enriched in the uropod (Figure 4A, pretubulysin). Colocalization of F-actin and pMyosin was found predominantly in the uropod and the perinuclear region (Figure 4A, pretubulysin). Quantification of our results shows that, while the level of F-actin was not affected drastically, pMyosin was significantly increased in pretubulysin-treated CTLs relative to their control counterparts (Figure 4B). Notably, colocalization between F-actin and pMyosin was also enhanced after pretubulysin treatment (Figure 4C). These findings indicate that disassembly of the MT network repolarizes the F-actin network and the associated actomyosin contractility to the rear part of CTLs.

**Figure 4.**
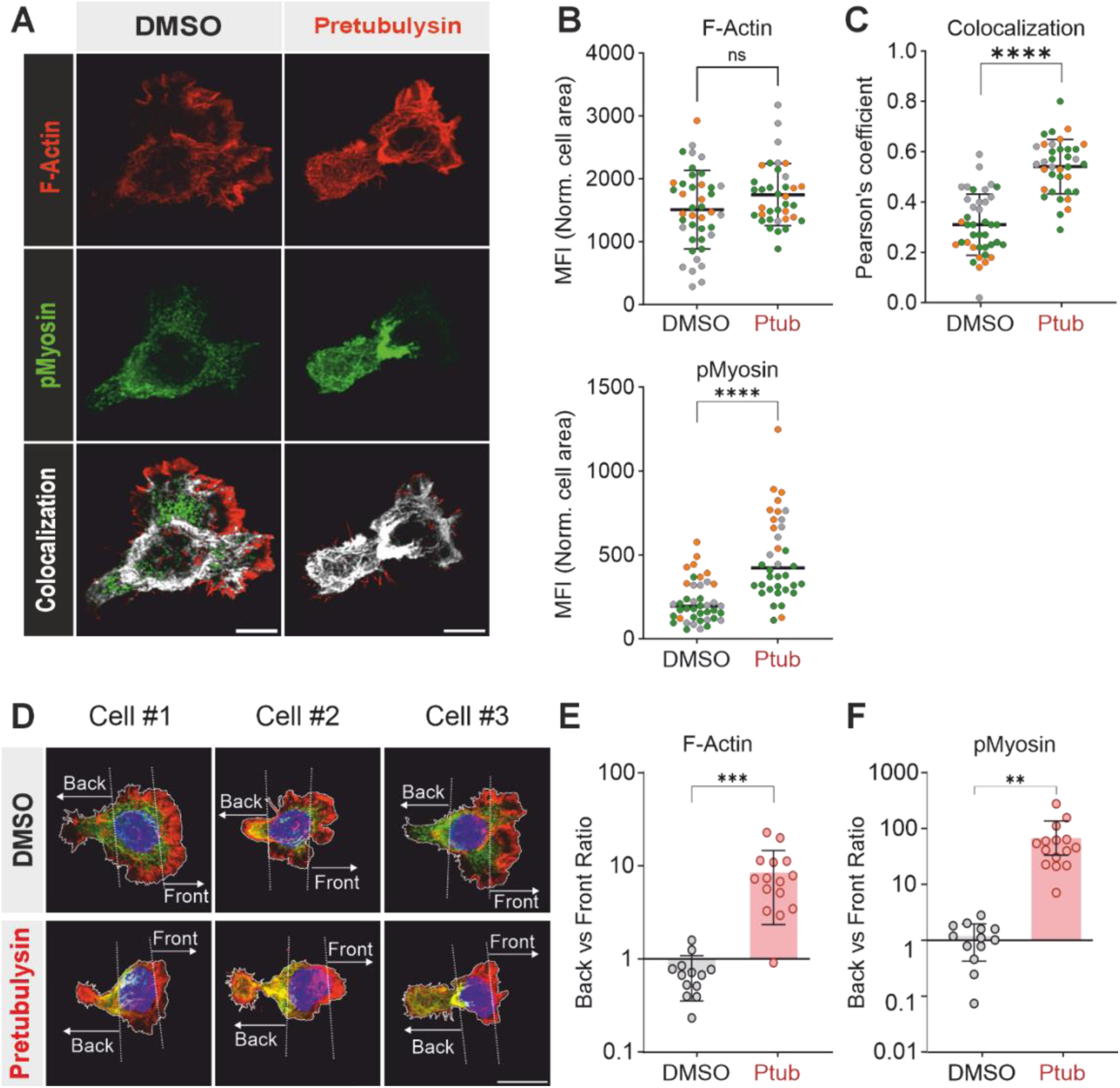
Pretubulysin leads to the reorganization of actomyosin. (A) Maximum intensity projection of representative CTLs treated with DMSO or pretubulysin (10 μM), immobilized on cell-tack coated coverslips, fixed/permeabilized, and immune-stained for F-actin (red), pMyosin (green) and the colocalized region of F-actin and pMyosin (grey). Colocalization analysis was carried out using ImageJ coloc2 plugin. Scale bar is 10 μm. (B) Quantification of F-Actin and pMyosin fluorescent signal on CTLs treated with DMSO or pretubulysin (10 μM). Cell border based on actin staining was defined as ROI. F-Actin and pMyosin in the region were quantified using Sum Intensity Projections with ImageJ and normalized with the cell area. Dots represent individual cells from three different donors including a color code for each donor. Error bars represent the standard deviation of the mean (mean ± SD). The Mann-Whitney test was used for statistical significance (**** p<0.0001). (C) Pearsons’s correlation coefficient was calculated using ImageJ coloc2 plugin. Dots represent individual cells. Error bars represent the standard deviation of the mean (mean ± SD). The Welch’s t-test was used for statistical comparison (**** p<0.0001). (D) Maximum intensity projection of three cells for each condition: DMSO and pretubulysin (10 M), immobilized on cell-tack coated coverslips. F-actin is shown in red, pMyosin in green and the nucleus in blue. The cell border (white) was established based on actin staining. “Back” and “front” regions were defined by the positioning of the nuclei and are represented with dotted lines (white) defined by the position of the nuclei. Scale bar is 10 µm. (E-F) Sum Intensity Projections were generated using ImageJ. Total F-Actin (E) and pMyosin (F) were calculated for “back” and “front” regions (MFI * region area). The graph represents the ratio between Front and Back. Dots represent individual cells and the mean ± SD is shown. The unpaired Student’s t-test with Welch’s correction was used for statistical significance (** p < 0.01, *** p < 0.001*).

While analyzing the images, we noticed that the morphology of pretubulysin-treated CTLs was also altered. To confirm this, we stained α-tubulin and nuclei in CTLs. In control cells, the MT network was clearly visible with a bright spot indicating the MTOC (SI Appendix, Fig. S7A, DMSO). In pretubulysin-treated CTLs, α-tubulin did not form filaments, was relatively evenly distributed in the cytosol, and a relatively bright spot, likely the MTOC, could still be identified (SI Appendix, Fig. S7A, pretubulysin). The orientation of CTLs was determined by MTOC positioning, which is always located at the uropod during migration. Interestingly, in MT-disassembled CTLs, the nucleus was relocated from its usual position at the front side edge (SI Appendix, Fig. S7A, pretubulysin). Quantifications of these results show that when the MT network was disassembled by pretubulysin, the uropod was bigger (SI Appendix, Fig. S7B) and the leading edge smaller (SI Appendix, Fig. S7C). The enlarged uropod might be a result of enrichment of actin and pMyosin, leading to enhanced contractile forces in this region.

To further characterize the relocation of actomyosin, we quantified the relative cellular distribution of F-actin and pMyosin. Fluorescence intensity was evaluated in two specific compartments: the back and the front of the cell. Three representative images are shown in Figure 4D for each DMSO and pretubulysin-treated CTLs. Quantification of the results again confirmed that F-actin and pMyosin are significantly accumulated at the back of pretubulysin-treated CTLs compared to the DMSO control, with an enhancement of around 10-fold for F-actin and around 100-fold for pMyosin (Figure 4E, F). Since actomyosin contractility plays an indispensable role in cell migration, we investigated whether the relocation of actomyosin to the uropod has any functional impact on migration. Thus, we studied the migration of CTLs upon MT destabilization and treatment with the ROCK-inhibitor Y27632. ROCK has been demonstrated to induce actomyosin-driven force bursts in migrating immune cells (38). We performed 2D migration experiments, in which we treated CTLs with DMSO, pretubulysin, pretubulysin and Y27632, or Y27632 alone. We used a 2D migration assay to be able to distinguish non-moving cells (with a track displacement < 10 µm) from moving cells. Pretubulysin induced a migratory phenotype, shown by an increased fraction of moving-cells and increased speeds (SI Appendix, Fig. S8A-C). In the combination treatment of pretubulysin and Y27632, we detected a significantly smaller portion of moving CTLs, reduced speeds, and a smaller mean-squared displacement (SI Appendix, Fig. S8A-C). We conclude that actomyosin machinery is required for the gained motility of CTLs by MT disassembly.

To gain further insight into the relationship between actomyosin distribution and cell migration behavior, we modeled amoeboid cell migration as a viscous droplet with an active boundary, which is analogous to the cell’s cortex. Relative cellular distribution of F-actin and pMyosin (back/front) is referred to as concentration ratio in this model. As shown in Figure S8D (SI Appendix), two cases with differing gradients of actomyosin concentration along the boundary were considered: Droplet 1 with low concentration ratio, representing DMSO-treated cells; and Droplet 2 with high concentration ratio, resembling pretubulysin-treated CTLs. The concentration profile is shown by the color scale (SI Appendix, Fig. S8D), where *c* is normalized by *c*_0_, the average droplet concentration. Both droplets were initialized with the same initial uniform concentration and ran until the droplet reached a steady state. The simulations show that for a motile active droplet, the translational velocity was dependent on the emergent gradient in boundary concentration of actomyosin: higher droplet speeds were obtained for droplets with greater concentration at the back of the cell, relative to the front of the cell (SI Appendix, Fig. S8E). These results show that enrichment of actomyosin leads to enhanced migration speed.

## Discussion

In summary, our work establishes pretubulysin as a potent MT disrupting agent that induces complete MT disassembly. Compared to nocodazole, pretubulysin treatment can further enhance CTLs migration, search and therefore killing efficiency in 3D collagen, particularly in dense collagen. Notably, MT-disassembled CTLs exhibit increased speed and persistence in microfabricated narrow channels, which mimic the confining tunnels in collagen matrices. This characteristic leads to overall enhanced motility and optimized searching efficiency in 3D environments as suggested by our random persistent searcher simulations. The role of actomyosin to create contractility to push cells forward has been shown in natural killer cells before (38). However, our observations go beyond that: we observed that MT disassembly results in elongation of the uropod and repolarization of F-actin and phosphorylated myosin from the protrusions at the leading edge towards the elongated uropod. This redistribution of the actomyosin network likely provides additional pushing forces for cell motility, favoring accelerated migration, as predicted by viscous droplet models, and shown in 2D migration assay. Our findings offer crucial insight into the intricate interplay between cytoskeletal dynamics and T cell function, offering potential avenues for enhancing immunotherapeutic strategies targeting T cell-mediated cytotoxicity.

The increased elimination of target cells in pretubulysin-treated CTLs despite unchanged killing capacity prompted the central mechanistic question of whether this enhancement can be explained fully by altered migration behavior, i.e., the experimentally observed increases in speed and persistence. Our PRW simulations were fed only with experimentally measured parameters (speed, persistence, and system geometry) with no fit parameters and showed quantitative, non-trivial agreement with experiments. This close agreement is not a trivial consequence of PRW dynamics but demonstrates that the observed killing enhancement can be explained entirely by altered migration behavior, without invoking changes in killing capacity or other mechanisms. It is also notable that while the effect of speed on the search efficiency is intuitive, the influence of persistence is not. The dependence of mean first-passage time on persistence is known to be nonmonotonic and system-size dependent (29). Thus, numerical evaluation has been necessary to predict the search time under observed experimental conditions. We emphasize that our conclusions are specific to this experimental context. More broadly, in addition to migration dynamics, killing capacity and other mechanisms may also contribute to the mechanistic basis of functional efficiency of immune cells. Another point is that our modeling approach simplifies the true geometry of the matrix and does not explicitly incorporate the channel network itself. In dense 3D collagen matrices, CTLs indeed migrate predominantly within narrow, pre-existing or self-generated channels, resulting in a highly guided and quasi-1D motility mode. This channel-like architecture leads to strong directional persistence, closely resembling the behavior observed in microchannel assays. For this reason, we considered speed and persistence values extracted from microchannel experiments to be appropriate inputs for our PRW model in the 3D collagen matrix. Indeed, the dynamics can be effectively decomposed into independent 1D components, and key transport quantities can be recovered via this decomposition. This offers a reasonable approximation for channel-guided movement in 3D matrices, if the matrix architecture does not introduce strong curvature or branching biases. Importantly, the close quantitative agreement between the model predictions and experimental killing data supports the plausibility of our assumption for the system under study.

A major advance of the present study is that it directly tests the requirement for microtubules in CTL cytotoxicity under conditions of significant MT disassembly. In contrast to previous studies, where residual microtubules could still support intracellular transport, pretubulysin treatment eliminated detectable MT filaments. Despite the disruption of the MT cytoskeleton, CTLs retained robust target-cell killing activity. To our knowledge, this is the first demonstration that efficient CTL-mediated cytotoxicity can occur in the absence of an organized MT network. These findings argue against a strict requirement for MT-dependent transport in lytic granule delivery and suggest that alternative, MT-independent mechanisms can support granule trafficking and secretion at the immunological synapse.

Beyond granule trafficking, microtubules are important regulators of CTL migration, which ultimately determines the efficiency with which CTLs search for and engage target cells. Previous studies have shown that partial MT disruption enhances CTL migration and killing in dense three-dimensional environments, although the underlying mechanism remained unclear (15, 18). Here, we provide mechanistic insight by showing that MT disassembly redistributes actomyosin from the leading edge to the uropod. This redistribution is expected to increase rearward contractility and generate greater propulsive forces, thereby accelerating cell migration. Notably, complete MT disassembly further increased migration speed compared with partial MT disruption, demonstrating a dose-dependent relationship between MT integrity and migratory behavior. While MT disassembly enhances CTL migration and cytotoxicity, intact microtubules may be maintained under physiological conditions to protect the nucleus and preserve cellular integrity during migration through mechanically restrictive environments.

In recent years, the MT network has been extensively studied as a therapeutic target for cytotoxic compounds in cancer. For the role of MTs in cell migration, it has been shown that destabilizing the MT network hinders mesenchymal migrating cells, such as fibroblasts or cancer cells, from generating cell protrusions in soft collagen gels (39). For endothelial cells, disruption of MTs inhibits trail retraction and therefore impairs cell migration on 2D surfaces and transmigration through membranes with pores (40). Immune cells mostly migrate through complex ECM environments in an adhesion-independent amoeboid manner, which is characterized by fast and adaptive shape changes (41). However, the migratory phenotype is versatile and depends on cell type and environment (41), thus also the role of MT destabilizing compounds in immune cell migration varies. *In vivo* studies in zebrafish have demonstrated that macrophages, Langerhans cells (skin-resident dendritic cells) and neutrophiles move upon MT destabilization, but the directionality and persistence are disturbed (42–44). In dendritic cells, MT dynamics are important for navigating the cells through pillar forests, by modulating retraction of protrusions via Rho A and its exchange factor Lfc (21). Interestingly, stabilization of the MT network with taxol perturbs cell migration as well (18, 45).

In summary, studies have demonstrated that MTs are important to maintain cell integrity and to guide the path for migrating immune cells (10). There seems to be a difference between larger and ramified cell types such as macrophages and dendritic cells, which require MTs to retract protrusions and find their path within complex environments (10) and smaller cells, such as T cells. When T cells were treated with MT destabilizing compounds, such as nocodazole, they move in a motile and persistent way (15, 18).

Thus, the MT network of T cells has emerged as an attractive target that can be specifically disrupted to maximize the efficacy of immune cell-related therapies. While MT depolymerizing agents have received considerable attention as cytotoxic drugs, there is a notable scarcity of studies examining their effects on T cells. Moreover, the existing studies predominantly focus on cancer cells rather than immune cells (46–48). Also, despite promising success, the therapeutic utility of many MT depolymerization agents is impeded by issues such as toxicity and resistance, prompting active exploration of novel compounds. Pretubulysin serves as a good example and is a synthetic precursor of tubulysin with accessible chemical synthesis while maintaining powerful antitumoral activity (24, 25, 49).

Our results show that persistence and speed are coordinated by the MT network in migrating CTLs. Our random persistence searcher simulations even suggest that persistence and speed can collectively tailor search and killing efficiency of CTLs in 3D environments. Besides their mechanical properties, MTs serve as a repository for guanine nucleotide exchange factors, which activate small GTPases that regulate actin assembly and actomyosin contractility (48, 50). Therefore, disruption of MTs can alter cell mechanics not only by changing the microtubule network mechanics, but also by inducing changes within the actin cytoskeleton (51). Previous studies reported that pharmacological dissociation of MTs leads to increased contractility (18, 20, 48). Furthermore, the interplay between microtubules and cellular contractility has been demonstrated to modulate the morphology and mechanical properties of migrating cells. The enhanced cell stiffness in MT-disassembled CTLs could be attributed to enhanced actomyosin contractility. Enriched actomyosin activity has been associated with cortex stiffening in non-adherent cells and monocytes (36). Although MTs do not directly contribute to the generation of forces that drive cell migration they are involved in the regulation of actin-dependent motility via guanine nucleotide exchange factor GEF-H1, in what has been recently described as the microtubule-contractility axis (18, 33). When GEF-H1 is bound to MTs, it is inactive. It becomes activated when MTs depolymerize, which happens either due to the inherent instability of MTs or due to the treatment with MT-depolymerizing compounds. Activated GEF-H1 activates Rho, which in turn induces the upregulation of myosin II contractility and actin polymerization (50–53). It is also reported that GEF-H1 plays an essential role in crosslinking the MT network and contractility in CTLs (18).

Actomyosin contractility provides primary driving forces for cells, including T cells, to move. To cross the barrier formed by the endothelium, T cells use actomyosin and the resulting contractility to squeeze between endothelial cells. The efficiency of migration is adjusted by the amount of force generated and the level of traction applied (54). While actin polymerization is essential for force generation by T cells, dynamic MTs at the interface also play a fundamental role (20). Actomyosin contractility at the uropod has been described as fundamental in generating the forces that drive migration in amoeboid cells (8). In this work, we observed that in pretubulysin-treated CTLs, F-actin and p-Myosin were redistributed from the protrusions to the uropod. The results from viscous droplet models suggest that enrichment of actomyosin at the uropod relative to the front of the cell can increase migration speed. This symmetry-break in the distribution of the cytoskeletal elements results in morphological changes, mechanical perturbations, and migration enhancement of CTLs, which can then kill their targets more efficiently.

## Materials and Methods

CTLs from healthy donors were pharmacologically perturbed and analyzed using 1D and 2D migration assays, 3D tumor killing assays and real-time deformability cytometry. Effects of MT-destabilizing compounds were further assessed using *in vitro* MT reconstitution in microfluidic devices and quantitative image analysis. We further complemented the experiments with computational simulations linking cell motility to search behavior, killing dynamics and actomyosin concentration in active droplet models. Full description of experimental procedures, reagents, imaging and analysis workflows, modeling, and statistical methods are provided in the SI Appendix, Materials and Methods.

## Acknowledgments

This work has been supported by funding from the German Research Foundation (DFG) CRC 1027 (to M.H., F.L., B.Q., L. Schaedel) and the NanoBioMed Young Investigator grant awarded by Saarland University to L.Schaedel.

