## Supplemental Information for "Revisiting the role of microtubules in cytotoxic T cell function: efficient search and killing without microtubule network"

**This PDF file includes:**

Supporting Information Text – Extended description of materials and methods
Figures S1 to S9
Legends for Movies S1 to S2
SI References

**Other supporting materials for this manuscript include the following:**

Movies S1 to S2

### Supporting Information Text

#### Materials and Methods

**Antibodies and Reagents.** The following antibodies were used: α-Tubulin (Invitrogen REF 32-2500), pMyosin light chain (Cell Signaling REF: 08/2019), Alexa Fluor 647 conjugated goat anti-rabbit (Invitrogen A21244), and Alexa 488 conjugated donkey anti mouse (Invitrogen A21202). The following reagents were used: Phalloidin-iFluor 594 (abcam REF: 176757) Hoechst 33342 (ThermoFisher Scientific), Fluoromount-G with DAPI (Invitrogen REF00-4959-52), Carboxyfluorescein succinimidyl ester (CFSE), FibriCol® type I collagen Solution (Bovine,

Advanced Biomatrix), cell-Tak (Corning), poly-dimethylsiloxane (PDMS) (RTV-615; Momentive Performance Materials), PEG-PLL (Susos, Dübendorf, Switzerland). The following drugs were used: pretubulysin (synthesized as described in (1), nocodazole (Sigma-Aldrich) and Y27632 (Cayman Chemical); The compounds were dissolved in DMSO at 10 mM and stored at −20 °C. For experiments, the compounds were diluted to 10 µM or 1 µM in cell culture medium. The ROCK-inhibitor Y27632 was diluted to 20 µM in cell culture medium.

**CTLs preparation and cell culture.** Human peripheral blood mononuclear cells (PBMCs) of healthy donors were isolated from the Leukocyte Reduction System Chamber using a gradient centrifugation (450 g, 30 min) with Lymphocyte Separation Medium 1077 (PromoCell). Remaining red blood cells were removed by the lysis buffer (155 mM NH4Cl, 10 mM KHCO3, 0.1 mM EDTA, pH=7.3). For migration and killing experiments, PBMCs were stimulated with Streptococcal Enterotoxin A (SEA) at 0.5 µg/ml for 30 minutes and then diluted 50X in AIMV medium (ThermoFisher Scientific) containing 10% FCS and Interleukin-2 (50 U/ml). After 5 days in culture, CTLs were isolated using Human CD8+ T Cell Isolation Kit (Miltenyi Biotec). All CD8+ T cells were cultured in AIMV medium (ThermoFisher Scientific) containing 10% FCS and recombinant human Interleukin-2 (ThermoFisher Scientific). NALM-6 pCasper cells were generated by Knörck et al. (2) and were cultured in RPMI-1640 (ThermoFisher Scientific) containing 10% FCS and 1% Penicillin-Streptomycin in the presence of puromycin (0.2 µg/ml).

**CTLs treatment with drugs.** Cells were treated with pretubulysin at 1 µM or 10, with nocodazole at 1 µM or 10 µM or with DMSO at 0.1 % for 30 minutes prior the experiments. For 1D migration experiments, cells were loaded in medium with the corresponding drug. For 3D migration, infiltration, and killing experiments, cells were washed after drug treatment, before being placed on top of collagen layer. For immunofluorescence microscopy, cells were fixed immediately after drug treatment (30 minutes). For RTDC experiment, treated cells were resuspended in cell carrier buffer with drugs at the corresponding concentration.

**Microfabrication of 1D channels.** Microchannels had a length of 800 µm and a squared channel width and height of 5 µm. Using nanoscribe-generated stamps, microchannels were fabricated by pouring polydimethylsiloxane (PDMS) precursor mixture (Sylgard 184, base:curing agent = 10:1) and curing at 70°C for 2 hours. Then, circular-shaped reservoirs (2mm) were drilled, and shape was adjusted using a razor blade. After the PDMS chips were brough into the correct size, they were cleaned by sonification for 30sec in 70% of ethanol. Chips were dried afterwards with air and activated for 30 sec at 300 mTorr by plasma treatment (air or oxygen). The binding was enhanced afterwards by placing the chips for 1 hour in the oven at 65 °C. For coating the microchannels, right before adding the coating solution (0.5 mg/mL PLL-PEG), chips were activated by air plasma at 300 mTorr for 1 min and after 30 minutes of incubation with the coating solution, chambers were washed with PBS.

**1D migration experiment.** The chambers prepared with PLL-PEG-coated channels were equilibrated for 1 hour at 37°C with cell culture medium either with pretubulysin 1 µM or 10 µM, nocodazole 1 µM or 10 µM, or DMSO 0.1%, before loading the cells. Cells were stained with Hoechst 33342 (200 ng/ml) for 30 minutes at 37°C and 5% $CO_2$. Treated CTLs were loaded in the chambers in 10 µl at 20 Mio cells/ml and the chip was covered with medium. We used a life cell epi-fluorescence microscope (Nikon) with a 10X objective lens (Plan-Neofluor, NA = 0.5) with temperature and $CO_2$ control (Live Cell Instrument, Korea). Cells migrated spontaneously for 15 hours and images were acquired every 3 minutes. Non-moving cells were filtered out by the design of the experiments, since only moving cells were able to enter the 1D channels.

**1D migration tracking and analysis.** Custom-written routines in Matlab (Mathworks, Natick, USA) were used for tracking analysis. First, images were rotated (bilinear interpolation) so that the channels are perfectly horizontal in the movies. To obtain a flatfield for correction of inhomogeneous background over the field of view (FOV), a large (30x30 pixels) median filter was applied to each fluorescence image. This removes all cells, leaving a flatfield, which was subsequently subtracted from each frame.

Positions of microfluidic channels were automatically identified. Positions of the microfluidic channels were identified as follows: First a maximum intensity projection over time was calculated, then all pixels were summed horizontally, leading to a peak for each channel that contained at least once a fluorescent cell. These peaks were detected with MATLAB's findpeaks command. Non-

channel regions were replaced by the average intensity of the image. The resulting images were smoothed with a Gaussian filter of 5 pixels full width at half maximum.
Cells were identified as local intensity maxima (Matlabs command imregionalmax) exceeding a threshold in the resulting images. Cells were subsequently tracked (i.e. re-identified in consecutive frames) by minimizing the squared distance between all particles in consecutive frames as described in (3). Each microfluidic channel was treated separately during tracking. Cell trajectories, interacting with or influencing other trajectories were excluded from the analysis. The quality of each track was controlled manually. Cell speeds were calculated as displacement between frames divided by the time interval between frames. Cell persistence was defined as the ratio of the net displacement (distance between last and first position) to the total displacement (length of path traveled by the cell). As an additional quality control measure, we examined track lengths to ensure that only actively migrating cells were included in the analysis. The track length distributions (Fig. S9) indicate that most trajectories span distances consistent with the channel length and the width of the field of view. Notably, CTLs treated with 1 µM nocodazole exhibited longer trajectories (Fig. S9), which is consistent with their reduced persistence observed in the motility analysis. These extended track lengths likely result from repeated back-and-forth movements within the channel.
**2D cell migration experiment.** Activated CLTs were treated for one hour with 10µM pretubulysin, 20 µM Y27632, a combination of both, or 0.3 % DMSO. In parallel cell nuclei were stained with 6.7 µM Hoechst 33342 trihydrochloride (Thermo Fisher, Germany) in each condition. After the treatment, $1x10^6$ cells per condition were centrifuged (300 g, 5 min) and resuspended in 200 µl medium containing the drugs. In order to avoid biased T cell movement due to convection and automated microscope stage movement the T cells were stabilized. For this, a two-component silicon glue (Picodent, Germany) was mixed in a 1:1 ratio and incubated at RT for 6 min. One drop of the glue was transferred to one edge of a well from 24 well plate and another silicon glue drop was transferred to the opposite sides of the same well. 10 µl of the cell suspension (corresponding to 50,000 cells) were transferred to the center of the well. A 12 mm round glass coverslip (Carl Roth GmbH + Co. KG, Germany) was layered on top adhering to the silicon glue and therefore fixing the T cells. The samples were incubated for 1 hour at 37 °C and 5 % $CO_2$ before tracking them for 15 hours every 2 min using a laser scanning confocal microscope LSM 880 (Carl Zeiss AG) equipped with a 10x objective (EC"Plan-Neofluar" 10x/0.30 M27, Carl Zeiss AG, Germany) and a 405 laser line (to visualize the Hoechst signal). Segmentation and tracking of migrating cells were based on the Hoechst signals. We tracked migrating cells for 120 frames (4 hours) using the Fiji (4) plugin TrackMate (5) with the built-in StarDist detector (6). Trajectories with track size < 50 frames and clusters of cells were removed. Trajectories with a track displacement below 10 µm were assigned as non-migrating cells and were also not considered for the analysis of the cell motility. Trajectories of migrating cells, consisting of the position (x,y) in each frame were analyzed using a custom-written MATLAB script. Instantaneous speed was calculated by dividing local displacement of the cell between two successive frames with the time interval (2 min). The mean squared displacement was calculated with the following formula:

$$MSD(\Delta t) = \langle [x(t+\Delta t) - x(t)]^2 + [y(t+\Delta t) - y(t)]^2 \rangle$$

**Preparation of Collagen Matrix.** As described in (7) bovine collagen type I stock solution was neutralized (pH 7.0-7.4) with 0.1 N NaOH and PBS 10X on ice. PBS was used to further dilute the collagen solution to the final experimental concentrations and distributed in the 96-well plates maintaining cool conditions. The collagen solution was finally incubated for 1 hour at 37°C with 5% $CO_2$ for fibrillation. We used collagen concentrations of 2 mg/ml and 4 mg/ml for the infiltration assay and 4 mg/ml for the killing assay.
**3D killing experiment.** For killing assays, NALM-6-pCaspe (NALM-6 cells expressing the apoptosis reporter pCasper-pMax) were used as target cells. NALM-6-pCasper were treated with staphylococcal enterotoxin A (0.1 µg/ml) for 40 min at 37°C with 5% $CO_2$, then resuspended in chilled collagen solution. Afterwards, we transferred them in 96-well plates, and centrifuged them at 4°C (200 g, 7.5 min) to sediment them on the bottom of the well. Afterwards, the mix target cells-collagen (50 µl/well) was solidified in the incubator for 1 hour at 37°C with 5% CO2 for collagen fibrillation. Afterwards, we added the CTLs on top of the solid collagen matrix in medium without drug or DMSO. The used ratio of effector (CTL) to target cells was 5:1. Images were taken by

ImageXpress (Molecular Devices) with Spectra X LED illumination (Lumencor) every 20min for 36 hours. Culture conditions were maintained at 37°C with 5% $CO_2$.

**3D infiltration experiment.** CTLs were stained with CFSE (5 µM in PBS + 4.5% FCS) at room temperature and protected from light for 15 min, washed once with PBS, then resuspended in culture medium AIMV +10% FCS, and kept at 37°C with 5% $CO_2$ for recovery during 1 hour. Then, CTLs were treated with pretubulysin 1 µM or 10 µM, nocodazole 1 µM or 10 µM or DMSO and loaded on top of solidified collagen matrix. Images were taken by ImageXpress (Molecular Devices) with Spectra X LED illumination (Lumencor) every 20 min for 24 hours. Culture conditions were maintained at 37°C with 5% $CO_2$.

**Degranulation assay.** To assess stimulation-induced degranulation, CTLs were incubated with Raji cells previously pulsed with staphylococcal enterotoxin A in the presence of BV421 anti-$CD107a^+$ antibody and protein transport inhibitor Golgi stop (BD Biosciences) at 37 °C with 5% $CO_2$ for 4 hours. Then, the cell suspension was stained with PerCP anti-human CD8 antibody for 30 minutes. The samples were analyzed using FACSVerse (BD Bioseceinces). The $CD8^+$ population was gated for CTLs. FlowJo v10 (FLOWJO, LLC) was used for data analysis.

**Immunostaining.** CTLs were immobilized on coverslips using the Cell-Tak adhesive (Corning) following the manufacturer instructions. Next, cells were added to the Cell-Tak treated coverslip, immediately after pretubulysin or DMSO treatment and incubated for 2 minutes. Cell-Tak treated coverslip were carefully washed with PBS. Right after, pre-warmed paraformaldehyde (PFA, 4%) was added and incubated for 10 minutes at room temperature. Next, coverslips with immobilized cells were carefully washed twice with PBS, permeabilized with TritonX-100 (0.05%) for 10 minutes and blocked with 2% BSA in PBS for 1 hour. Staining with the indicated antibody or Phalloidin was performed in PBS+BSA 2% for time and dilution indicated in the antibodies and reagents section. DAPI was added to the slides for nuclei staining and coverslips were placed on mounting slides for imaging. Fixed/stained cells were imaged using the 63x immersion oil objective (Zeiss, Plan-Apochromat 63x/1.40 oil DIC M27) of a Zeiss LSM 900 confocal microscope with the Axiocam 705 Mono camera (Zeiss).

**Real-time deformability cytometry (RT-DC).** CTLs were treated with pretubulysin or DMSO for 30 minutes, pelleted by centrifugation and resuspended in 100 µl of Cell Carrier B solution (phosphate-buffered saline + long-chain methylcellulose polymers of 0.6 % w/v) with pretubulysin or DMSO. A 20 µm microfluidic PDMS chip was assembled on the stage of an inverted microscope (Zeiss). CTLs homogenously resuspended were loaded on the chip using a syringe pump. Using a CMOS camera, CTLs were live imaged while flowed through the channel were. At least 3000 events were acquired for each condition (pretubulysin or DMSO treated) and experiment, flowrate (0.04 to 0.12 µl/s) was used, according to the range suggested by the manufacturer for the channel size and carrier buffer. The stiffness of the cells was analyzed using ShapeOut (Zell Mechanik, Dresden). We used linear mixed models provided by the manufacturer to calculate statistical significances.

**Quantification of fluorescence intensity and colocalization analysis**

Quantification of fluorescence intensity and colocalization was done with Fiji (4). For each cell, a per-cell ROI was defined manually using the actin channel (Channel 2). Cell boundaries were delineated using the Wand (tracing) tool to encompass the full actin cytoskeleton, including protrusions. ROIs were added to the ROI Manager and applied uniformly across all Z-slices of the stack. The same ROI definition strategy and parameters were used for control and treated cells. Mean fluorescence intensities of myosin and actin were quantified within the per-cell ROI Measurements were restricted to pixels inside the ROI to exclude background and empty image regions. Identical measurement settings were applied to all images. Colocalization between myosin (Channel 1) and actin channel (Channel 2) was assessed using the Coloc 2 plugin in Fiji (4). Analysis was performed on full 3D stacks, with voxel-by-voxel comparison across all Z-slices within the per-cell ROI. Pearson's correlation coefficient was extracted as a measure of pixel-wise intensity correlations between channels.

**In vitro reconstitution assays. Tubulin purification and labeling:** Tubulin was purified from fresh bovine brains by three cycles of assembly and disassembly in high salt buffer [High Molarity PIPES Buffer: 1 M PIPES, pH 6.9, supplemented with KOH, 10 mM $MgCl_2$, 20 mM EGTA]. Use of high salt ensured that the purified tubulin was free of contamination from microtubule associated proteins as described in (8). We then labeled the purified tubulin with biotin or fluorescent dyes -

ATTO488 (ATTO-TEC, AD488) and ATTO565 (ATTO-TEC, AD565) dyes as described in (9). Silane-PEG-Biotin passivated cover glasses and ATTO-565 labeled-biotinylated microtubule seeds were prepared according to protocol described in (9).

**Microfluidic Circuit Fabrication:** Using standard soft lithography, the microfluidic chip with an inlet and outlet port was fabricated using PDMS (Sylgard 184, Dow Corning). TFE Teflon tubing (Supelco, inner diameter: 0.8 mm, outer diameter: 1.58 mm, Merck) was inserted into the outlet port. Tubing with an inner diameter of 0.03 mm and outer diameter of 1.58 mm was used to connect the inlet with the sample reservoir, via a manual shut-off valve connected to a pressure controlled microfluidic pump (LineUP Flow EZ 345 mbar, Fluigent). Refer Fig. 3A for the experimental setup.

***In vitro* reconstitution experiments using microfluidics:** For *in vitro* assays, 10 mM stock solutions of pretubulysin and nocodazole (dissolved in DMSO) were further diluted in 1xBRB80 [Brinkley Buffer 80: 80 mM PIPES, pH 6.8, 1 mM EGTA and 1 mM $MgCl_2$] buffer. The PDMS chip was mounted onto a passivated cover glass and fixed on to the microscope stage. The chip was first equilibrated with a solution of 1xBRB80. The surface was then perfused with 300 µl of Neutravidin (50 µg $µl^{-1}$ in BRB80; Pierce), followed by 300 µl of PLL-g-PEG (Pll 20K-G35-PEG2K, Jenkam Technology) at 0.1 mg/ml in 10 mM Na-HEPES buffer (pH 7.4) followed by another wash with 1xBRB80. ATTO-565 (Red) labeled biotinylated microtubule seeds were then flushed into the chamber and allowed to bind for a period of 5 mins. Unbound seeds were then removed by subsequent washes with BRB80 supplemented with 1% BSA. Microtubules were polymerized from the attached seeds by addition of elongation mix (56 mM PIPES, 0.7 mM EGTA, 0.7 mM $MgCl_2$, 38 mM KCl, 19 mM Phosphate buffer, pH 6.8) containing 10 µM of tubulin (20 % ATTO-488 labeled), supplemented with 1 mM GTP, an oxygen scavenger cocktail (20 mM dithiothreitol, 1.2 mg $ml^{-1}$ glucose, 8 µg $ml^{-1}$ catalase and 40 µg $ml^{-1}$ glucose oxidase), 1 mg/ml BSA and 0.025 % methyl cellulose (1500 cp, Sigma). After 10 min of elongation, the drug containing mix [elongation mix containing 10 µM of tubulin (to prevent microtubule disassembly due to dilution) along with various concentrations of the drugs (pretubulysin/nocodazole)] was perfused into the chamber. For control experiments, elongation mix containing 10 µM of tubulin with equivalent concentrations of DMSO was flushed in. Microtubules were imaged before, during and after addition of each drug.

**Imaging and analysis:** MTs were visualized using a 63x oil immersion objective (Zeiss, Plan-Apochromat 63x/1.40 oil DIC M27) on a Zeiss LSM 900 confocal microscope with the Axiocam 705 Mono camera (Zeiss). Experiments were performed at 37° C using a stage controller (Insert-P, PeCon). Time-lapse recording (with a frame interval of 1s) of both 488 and 565 channels (using 0.08 mW laser output power) was performed using the line scan mode in the Zen blue software (version 3.2, Zeiss). We used the subtract background and smooth functions of Fiji (version 1.53t) to increase the signal/noise ratio in our videos (4). MT mass before and after addition of the drug was calculated by measuring the total length of microtubules in a single field of view over a period of 8 min.

**Persistent random search simulations.** We performed Monte Carlo simulations of persistent random walks in a three-dimensional box of lateral size L × L and height h, chosen to mimic the geometry of the experimental collagen matrices. Unless stated otherwise, we used L = 5.6 mm and h = 1.5 mm. The simulation volume (Figure 2A inset) was laterally confined by reflective boundaries. In the vertical direction, walkers moved between a reflecting entry plane at the top and an absorbing plane at the bottom where target cells were located. No directional bias or external force was applied; all walkers performed an unbiased persistent random walk.

Each simulated CTL started at a random position on the top plane with a random incident direction toward the interior. Walkers were allowed to move back toward the entry plane during the simulation but could not exit through it. A trajectory terminated once the walker reached a target on the bottom plane. Targets were represented as non-overlapping disks of area ~200 µm², randomly distributed over the bottom plate (~25,000 targets in total). Upon first contact, the target was removed to mimic immediate elimination after engagement. Killing events were monitored inside a central observation window of 0.7 mm², consistent with the experimental field of view. For the comparison of numerical and experimental covered area in Fig.2A, the target layer in simulations was considered as a discrete 2D grid that was initially fully occupied.

Simulations progressed in discrete time steps of $\Delta t$ = 30 s. At each step, the direction of motion was updated according to the standard persistent random walk scheme: the polar angle change $\theta$ relative to the previous direction was sampled from $\theta = \arccos(p)$ (10, 11), where p is the persistence parameter; the azimuthal angle was chosen uniformly from [0,2π]. Instantaneous speeds were drawn from an exponential distribution with the experimentally measured mean speed. In earlier work, this framework was extended to multistate dynamics (12, 13) to describe short-term motility modes in collagen (14); however, because the present study focuses on timescales well beyond the transient regime, we employed a single-state persistent random walk model.

To enable direct comparison to experiments, all simulation parameters (mean speed, persistence, and geometry) were taken from independent measurements. In particular, speed and persistence of control CTLs were taken from 3D collagen migration data, whereas parameters for pretubulysin-treated CTLs were estimated from fold-changes measured in the microchannel assays. Although channel-guided migration in collagen is not explicitly modeled, the effective persistent random walk approximation is validated by the quantitative agreement between simulated and experimental killing dynamics (Figure 2A), supporting its suitability for the present system.

**Active droplet simulations.** The active droplet is modelled as a viscous fluid with an active boundary, implemented using the immersed boundary method. Full details of the numerical implementation of the hydrodynamics and immersed boundary can be found in (15). The active boundary of the droplet has an associated concentration representing active contractile particles which spontaneously breaks symmetry, leading to gradients in boundary tension driving droplet motion. The droplet is initialized with a uniform concentration on the boundary $c_0$ and simulations are run to steady state. To tune the concentration profile to values similar to those seen in experiment we set a concentration maximum by introducing a tangential forcing term on the boundary above the threshold concentration. We set the concentration threshold to [1.5 $c_0$, 2,5 $c_0$] for the low and high case respectively.

**Statistical Analysis.** For RT-DC, linear mixed models are included in the ShapeOut software. GraphPad Prism 9.5.1 Software (GraphPad) was used for statistical analysis of the rest of the experiments. In Graph Pad, first normality was tested (D'Agostino and Pearson). We compared two groups, which were normally distributed with paired t-test. If there was no normal distribution, the Mann–Whitney–U-test was used. If more than two groups were compared, we used a one-way ANOVA test (Kruskal Wallis test, if data was not normally distributed) for statistical comparison and Dunn's test to compare each group with each other. Statistical significance was defined as follows: * $p < 0.05$, ** $p < 0.01$, *** $p < 0.001$, **** $p < 0.0001$. Datapoints from different donors per condition were pooled for statistical testing.

**Ethical Considerations.** Our work for this study with healthy donor material (leukocyte reduction system chambers from human blood donors) was authorized by the local ethic committee [declaration from 16.4.2015 (Ha 84/15; Prof. Dr. Rettig-Stürmer) and amendment from 23.03.2021 (Ha 84/15; Prof. Dr. Markus Hoth)].

**Supplementary figures**

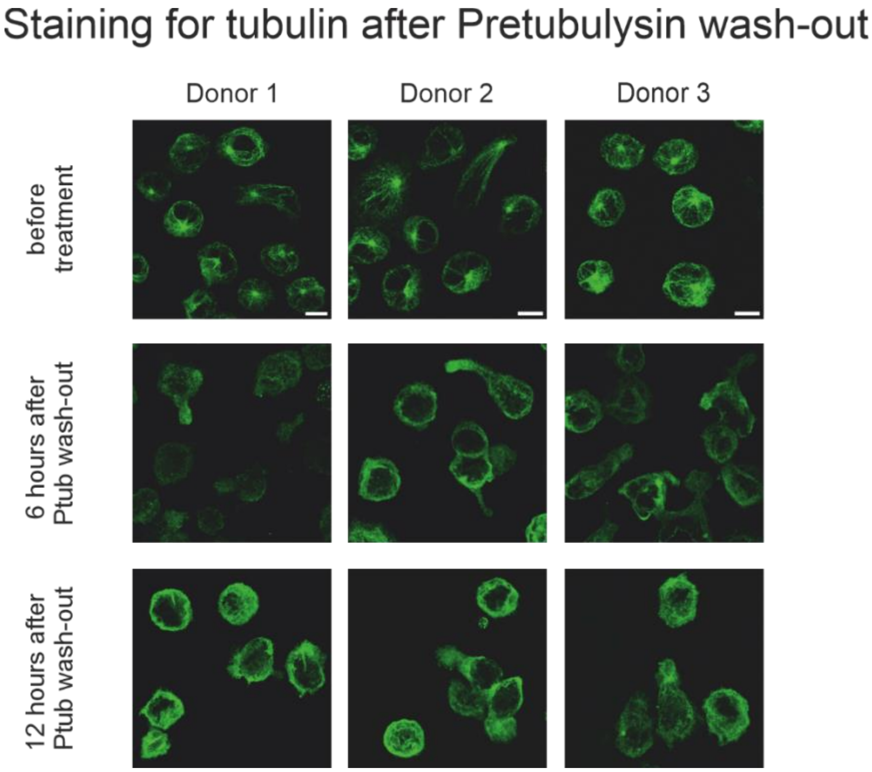


**Fig. S1. MTs remain perturbed after pretubulysin wash-out.** CTLs were fixed and stained for tubulin before treatment (row 1), 6 hours (row 2) and 12 hours (row 3) after pretubulysin treatment with a subsequent wash-out. Representative images show the MT network for the different conditions of three independent donors. Scale bar is 10 µm.

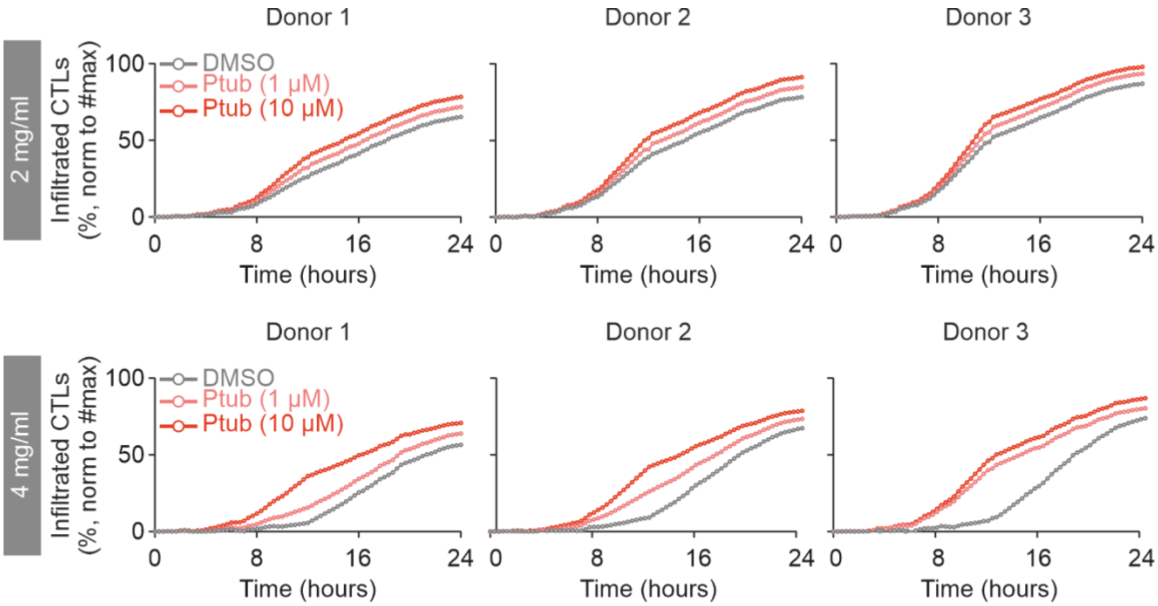


**Fig. S2. Time-resolved CTL infiltration into collagen matrices following pretubulysin treatment.** Time-resolved infiltration of cytotoxic T lymphocytes (CTLs) into collagen matrices of two different densities (2 and 4 mg/mL). CTLs were labelled with carboxyfluorescein succinimidyl ester (CFSE), treated with DMSO or pretubulysin (Ptub; 1 or 10 μM), and subsequently added to the top of polymerised collagen matrices. Cells reaching the focal plane at the bottom of the collagen matrix were quantified over 24 h, with images acquired every 20 min. Data are shown separately for three independent donors (Donors 1–3). Two technical replicates were performed for each donor and experimental condition and averaged at each time point. The number of infiltrated CTLs was normalised to the single maximum number of infiltrated CTLs.

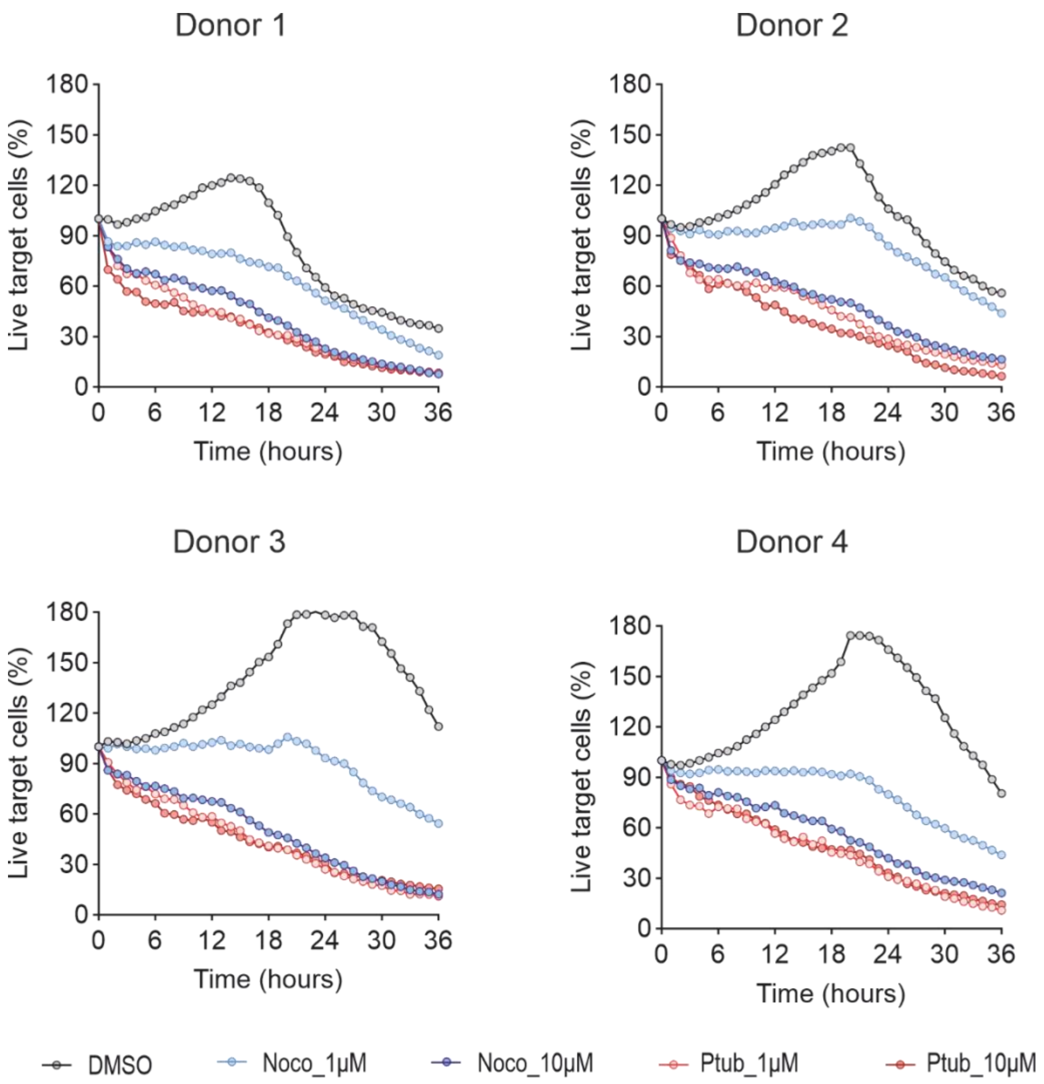


**Fig. S3. Results of the killing assay for each individual donor.** Figure shows the killing assay per treatment condition for each donor individually. The data from all four donors together is shown in Figure 1C.

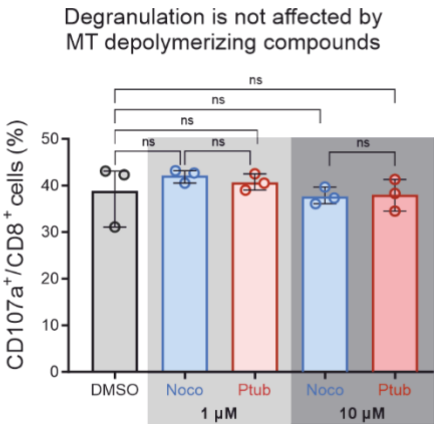


**Fig. S4. CTL's degranulation capacity is not affected by microtubule depolymerization with pretubulysin or nocodazole.** To determine the degranulation of CTLs after treatment with MT destabilizing agents, the fraction of $CD107a^+$ expressing CTLs over all activated CTLs ($CD8^+$) was determined using flow cytometry. Means of three biological experiments were compared using Kruskal-Wallis ANOVA with Dunn's multiple comparison test.

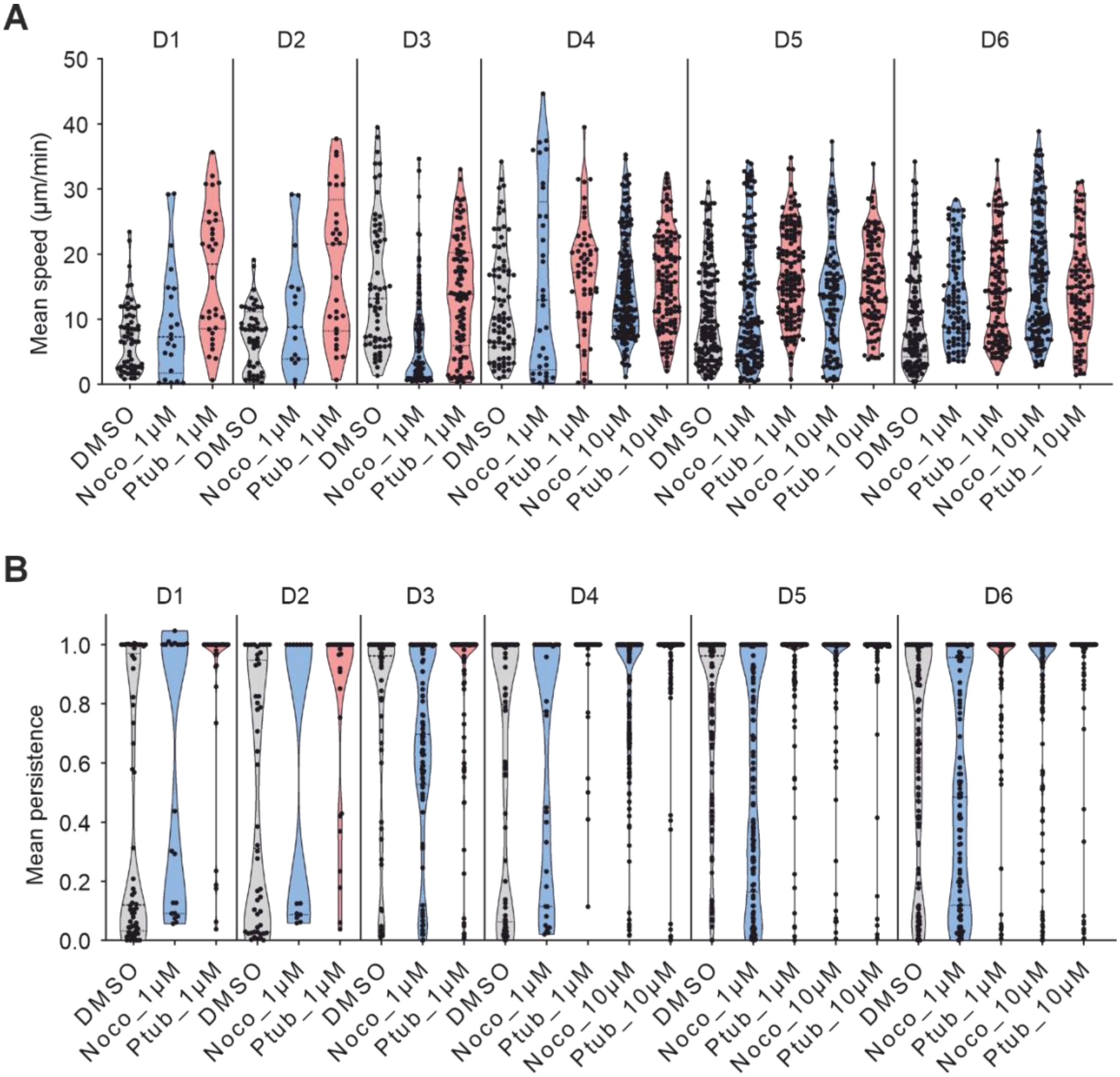


**Fig. S5. 1D Migration results for each donor.** Representation of mean speed (A) and mean persistence (B) for each individual donor. Six donors were tested for DMSO and low concentrations of nocodazole and pretubulysin (1 µM), while three donors were assessed for the higher concentrations (10 µM). The data from all donors together is shown in Figure 1E and F.

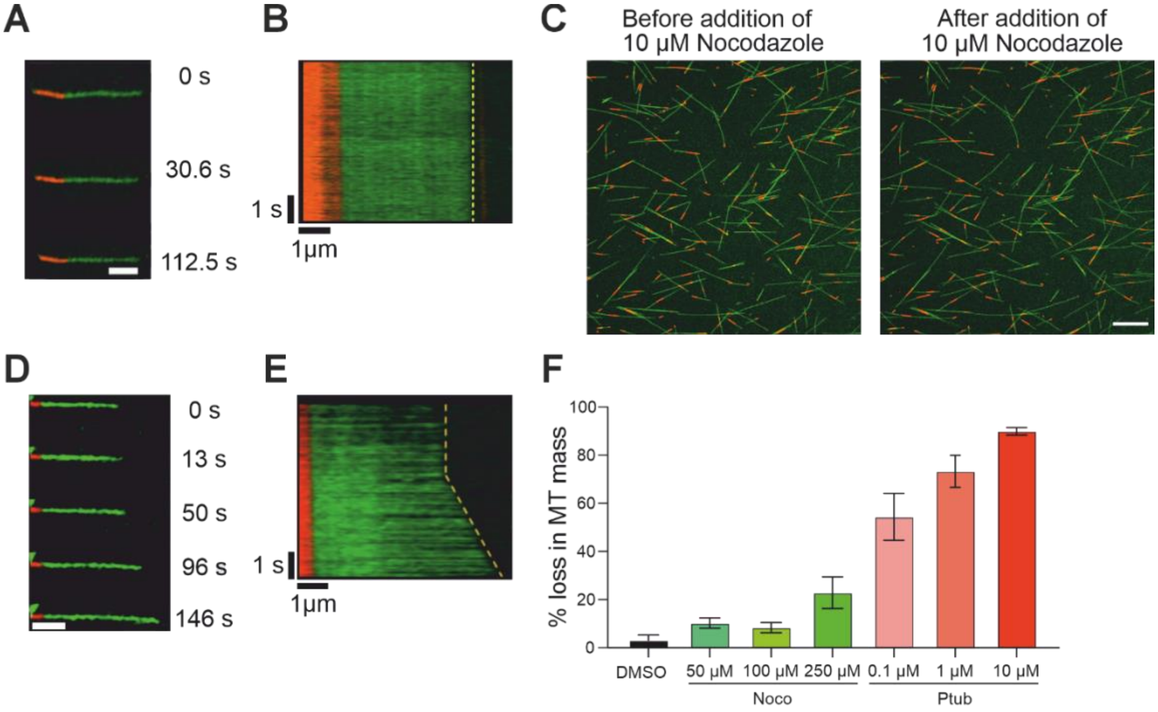


| | Control (DMSO) | Pretubulysin 0.1 μM | Pretubulysin 1 μM | Pretubulysin 10 μM |
|---|---|---|---|---|
| **Control (DMSO)** | - | 0.0025 | <0.0001 | <0.0001 |
| **Nocodazole (50 μM)** | 0.0158 | 0.0038 | <0.0001 | <0.0001 |
| **Nocodazole (100 μM)** | 0.0414 | 0.0034 | <0.0001 | <0.0001 |
| **Nocodazole (250 μM)** | 0.0079 | 0.0211 | 0.0007 | <0.0001 |

**Adjusted p-value Tukey multiple comparison test**

**Fig. S6. Characterizing microtubule depolymerization potential of nocodazole using *in vitro* reconstitution assays**

(A) Timelapse sequence showing microtubule pause (with no subsequent change in microtubule length) following treatment with 10 μM nocodazole (Scale bar 2 μm). Images are representative of three independent experiments. (B) Kymograph from A, depicting no change in microtubule length vs time after treatment with 10 μM nocodazole. (C) Representative image of a field-of-view before and after addition of 10 μM nocodazole (Scale bar 10 μm). Images are representative of three independent experiments. (D) Timelapse sequence showing microtubule pause and then slight elongation following treatment with 10 μM nocodazole (Scale bar-2 μm). Images are representative of three independent experiments. (E) Kymograph from D, depicting pause and increase in microtubule length vs time after treatment with 10 μM nocodazole. (F) Comparison of % loss in microtubule mass following treatment with nocodazole (50, 100 and 250 μM), pretubulysin (0.1, 1 and 10 μM) and Control (DMSO). Data represent Mean ± SD from three independent experiments. Unpaired t-test was used for statistical significance. Refer to the table for the corresponding p-values.

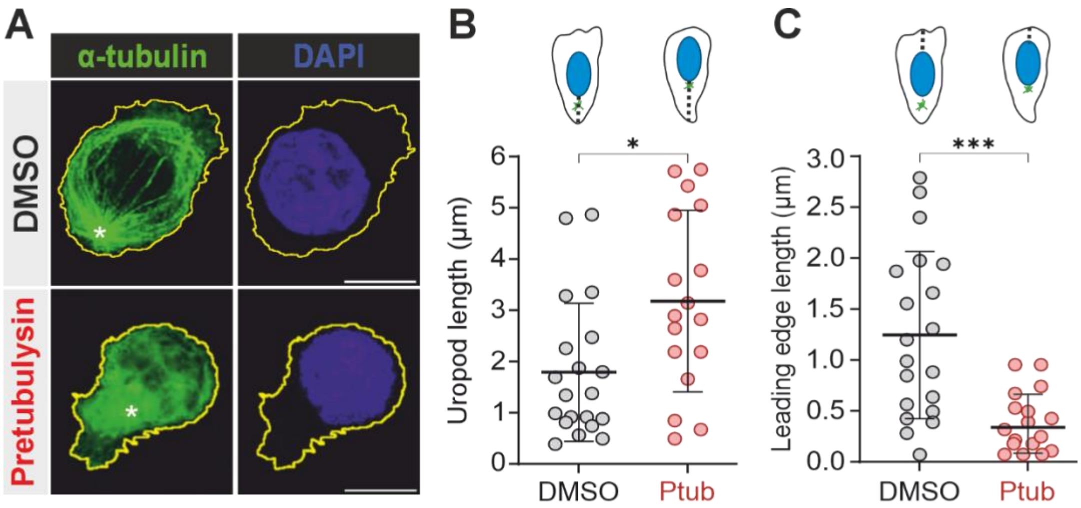


**Fig. S7. Microtubule disruption by pretubulysin changes morphology of CTLs.**

(A) Maximal Intensity projection of one representative cell for each condition (DMSO and pretubulysin 10 μM), immobilized on cell-tack coated coverslips showing that pretubulysin induced MT network disassembly on CTLs. Cell border based on actin staining is shown in yellow, α-tubulin in green, and nucleus in blue. The arrows indicate the MTOC. Scale bar is 10 μm. (B-C) Length of the uropod and of the leading edge were calculated manually using ImageJ from fluorescent confocal images. Schematics at the top represent CTLs under two conditions: DMSO or pretubulysin (10 μM) treated. Nucleus is blue, the MTOC is green, and the dotted line represents the distance measured. For uropod, distance was from the cell edge to the nuclei. For leading edge, distance was from the nuclei to the cell edge. On the graphs, dots represent individual cells from two donors. Error bars represent the standard deviation of the mean (mean ± SD). For statistical significance, the Mann-Whitney test was used for analyzing the back and front distance (* $p < 0.05$ and **** $p < 0.0001$, respectively).

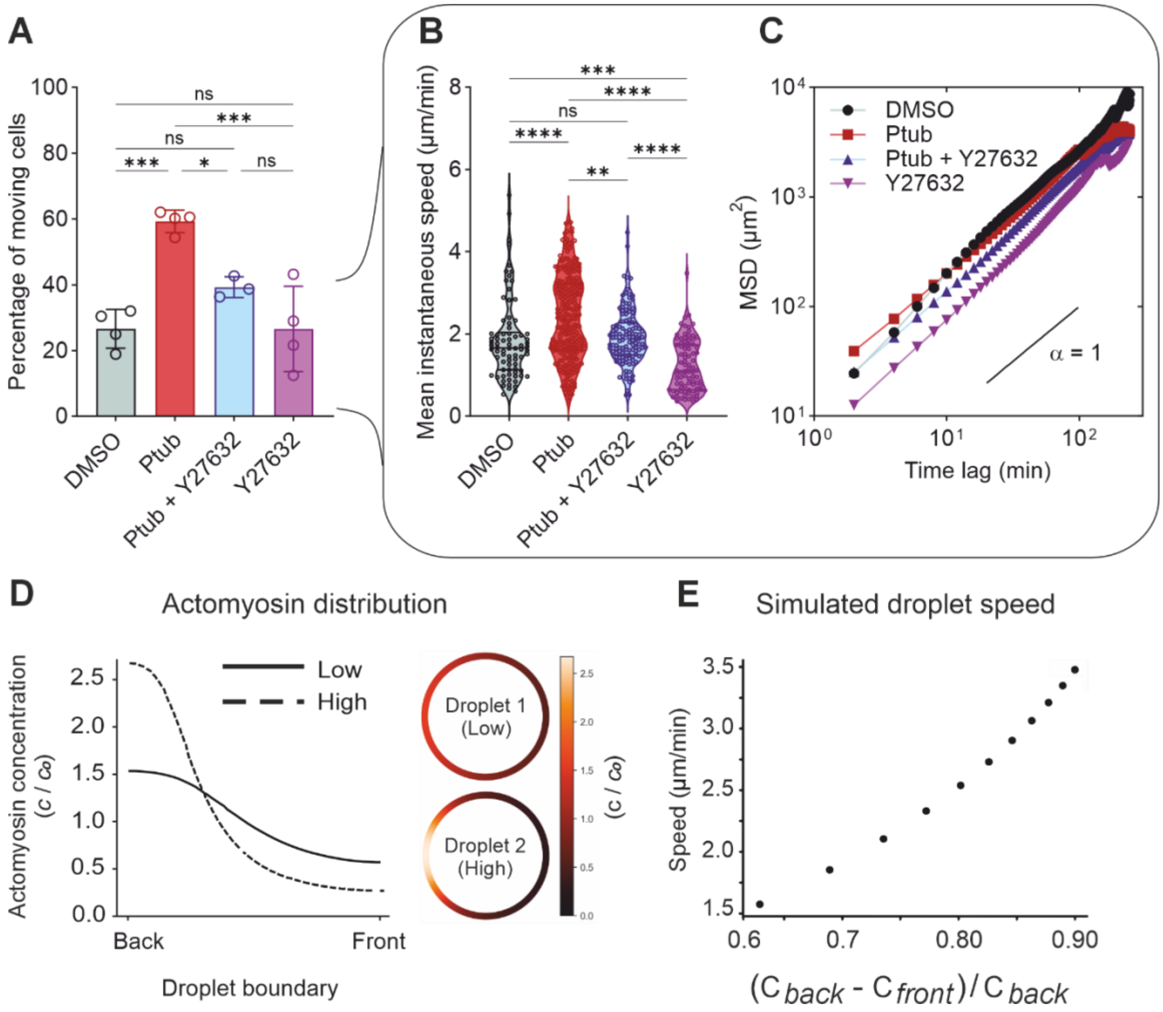


**Fig. S8. Microtubule disruption-induced actomyosin accumulation at the uropod favors migration.**

CTLs were treated with DMSO (grey), pretubulysin (red), a combination of pretubulysin and the ROCK-inhibitor Y27632 (blue), and Y27632 (violet) alone. Movements were tracked in 2D with live cell microscopy. Trajectories were assigned as moving cells, once track displacement was > 10 μm. The fractions of moving cells per conditions for four field of views (three field of views in Y27632 + Ptub) are shown in (A) and were compared using ANOVA with Dunn's multiple comparison test (* $p < 0.05$, *** $p < 0.001$). (B) Mean instantaneous speeds from trajectories, which have been assigned as moving, were compared using Kruskal-Wallis test with the Dunn's multiple comparison test (** $p < 0.01$, *** $p < 0.001$, **** $p<0.0001$). (C) The resulting mean squared displacements from moving cells in each condition. (D-E) Using computational hydrodynamics amoeboid cell migration was modeled as a viscous droplet with an active boundary analogous to the cell's cortex. (D) Emergent gradients in concentration of actomyosin were tuned to reflect experimental profiles. Droplet 1 represents an example of a small difference between back and front (low ratio), comparable with the experimental data obtained for control (DMSO) CTLs. Droplet 2 represents an extreme example of high difference between back and front (high ratio) comparable with the experimental data obtained for pretubulysin (10 μM) treated CTLs. The concentration profile is shown by the color scale, where $c$ is normalized by $c_0$, the average droplet concentration. Both droplets have equal total and average concentration. (E) Droplet speed against concentration where the difference was normalized against the concentration at the back of the droplet. The calculations indicate that greater difference in actomyosin from front to back leads to faster migration.

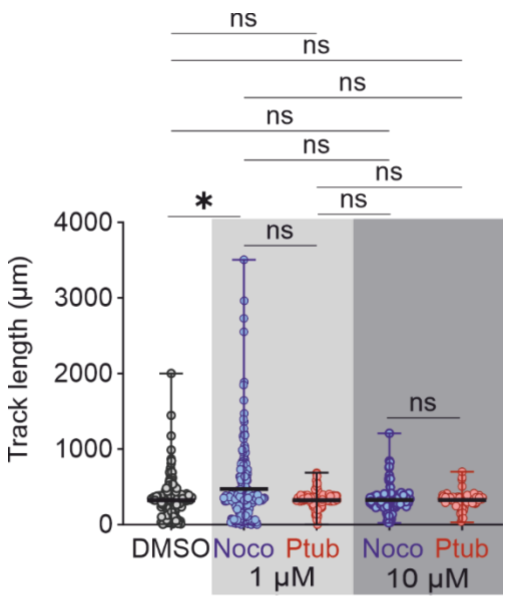


**Fig. S9. Track lengths of trajectories from 1D migration experiments.** Mean track lengths were compared using Kruskal-Wallis ANOVA with Dunn's multiple comparison test (* $p < 0.05$).

**Supplementary movie legends**

**Movie S1: Time lapse movies of infiltrating CTLs.** Time-resolved infiltration of CTLs into collagen matrices of two different densities (2 and 4 mg/ml). CTLs were labeled with CSFE, treated with DMSO or pretubulysin (Ptub 1 or 10 μM), and subsequently added to the top of polymerized collagen matrices. Cells reaching the focal plane at the bottom of the collagen matrix were quantified over 24 hours, with images acquired every 20 minutes for donors and two technical replicates were performed for each donor. Results from Donor 1 is shown here. Scale bars = 50 μm.

**Movie S2. Time lapse movies showing tumor cells expressing pCaspar to visualize living (orange-yellow), apoptotic (green) and dead cells (lose fluorescence).** CTLs encounter and kill target cells after navigating through the collagen gel. Images were acquired every 10 minutes for donors and four technical replicates were performed for each donor. Results from Donor 1 is shown here. Scale bars = 50 μm..
